# Host-derived oxidized phospholipids initiate effector-triggered immunity fostering lethality upon microbial encounter

**DOI:** 10.1101/2023.11.21.568047

**Authors:** Marco Di Gioia, Valentina Poli, Piao J Tan, Roberto Spreafico, Anne Chu, Alex G Cuenca, Philip LSM Gordts, Laura Pandolfi, Federica Meloni, Joseph L Witztum, Janet Chou, James R Springstead, Ivan Zanoni

## Abstract

Macrophages detect invading microorganisms via pattern recognition receptors that recognize pathogen-associated molecular patterns, or via sensing the activity of virulence factors that initiates effector-triggered immunity (ETI). Tissue damage that follows pathogen encounter leads to the release of host-derived factors that participate to inflammation. How these *self*-derived molecules are sensed by macrophages and their impact on immunity remain poorly understood. Here we demonstrate that, in mice and humans, host-derived oxidized phospholipids (oxPLs) are formed upon microbial encounter. oxPL blockade restricts inflammation and prevents the death of the host, without affecting pathogen burden. Mechanistically, oxPLs bind and inhibit AKT, a master regulator of immunity and metabolism. AKT inhibition potentiates the methionine cycle, and epigenetically dampens *Il10*, a pluripotent anti-inflammatory cytokine. Overall, we found that host-derived inflammatory cues act as “*self*” virulence factors that initiate ETI and that their activity can be targeted to protect the host against excessive inflammation upon microbial encounter.

## Introduction

The immune system evolved to protect the host against infections and to maintain tissue and organismal homeostasis. This is accomplished through a delicate balance between immune resistance, tissue tolerance and damage, and repair. Phagocytes of the innate immune system initiate the immune response to pathogens by sensing pathogen-associated molecular patterns (PAMPs) via pattern recognition receptors [1]. Phagocytes also detect the presence of host-derived damage-associated molecular patterns (DAMPs) that are released by the host under stress conditions. DAMPs are major drivers of sterile inflammatory diseases and, similarly to PAMPs, are widely believed to be recognized via PRRs [2]. Since the “infectious *non-self*, non-infectious *self*” theory [3] and the “danger model” [4] were proposed, much discussion raised about the distinct contribution of PAMPs and DAMPs to the immune response elicited against pathogens [5]. For example, inflammasome activation plays key anti-microbial roles, and it has been associated with the sensing of both *self* and non-*self* signals [6]. Nevertheless, so far, no consensus has been reached on whether the DAMPs that activate the inflammasome, such as ATP or uric acid crystals, are necessary to control pathogen growth.

Beside utilizing PRRs, phagocytes can also recognize the presence of invading microorganisms by sensing changes in physiological processes caused by virulence factors released by pathogens. The inhibition of critical immune pathways by the activity of virulence factors initiates an effector-triggered immunity (ETI) that does not rely on PRRs. While this mode of activating the immune response is well-described in plants and is known as the “guard model” [7], ETI has been reported also in eukaryotic cells [8–10]. For example, the ICP0 viral effector initiates ETI by degrading MORC3, a host repressor of viral replication. MORC3 degradation in turn unleashes the induction of anti-viral interferons, independently of PRR stimulation [11]. So far, no example of DAMPs inducing an ETI response has been described.

Our work centers on a set of DAMPs derived from non-enzymatic oxidation of phosphocholine-containing phospholipids (hereafter oxPLs). While oxPLs drive inflammatory diseases such as atherosclerosis and nonalcoholic steatohepatitis [12, 13], their role during an infection is controversial. oxPLs have been shown to increase inflammation during lung virus infections via activating the PRR Toll-like receptor 4 (TLR4) [14, 15], while their role during bacterial encounter remains debated [16]. Administering exogenous oxPLs before exposure to bacteria reduces inflammation [17]. In contrast, when oxPLs are administered after exposure to PAMPs, they increase the production of pro-interleukin (IL)-1β and trigger the inflammasome activation [18–20]. Of note, the capacity of the host to produce oxPLs and their relevance in shaping the immune response during microbial encounter remain unknown.

Here, we tested whether oxPLs are produced upon microbial encounter and whether they contribute to the lethal inflammatory response that follows: i) a polymicrobial systemic infection; ii) the encounter of methicillin-resistant *S. aureus* (MRSA); and iii) a viral mimic that causes lung tissue damage.

## Results

### oxPLs are induced and increase mortality and morbidity following microbial detection

oxPLs are DAMPs that react with a wide variety of innate PRRs [16] and antibodies, including the innate natural IgM antibody E06 [21], which can be used to assess oxPLs levels *in vivo* [12, 13]. We initially tested whether oxPLs are increased during the immune response initiated by PRR stimulation. The levels of circulating oxPLs, as well as their accumulation in the spleen, were tested in mice injected systemically with distinct classes of PAMPs. We found that PRR stimulation induced the accumulation of oxPLs in the blood and in the white and red pulp of the spleen (**Figure 1A, B,** and **Figure S1A**). We also found that among the different immune cells that we tested (**Figure S1B**), macrophages showed the most significant accumulation of oxPLs (**Figure 1C, Figure S1C**).

**Figure 1:**
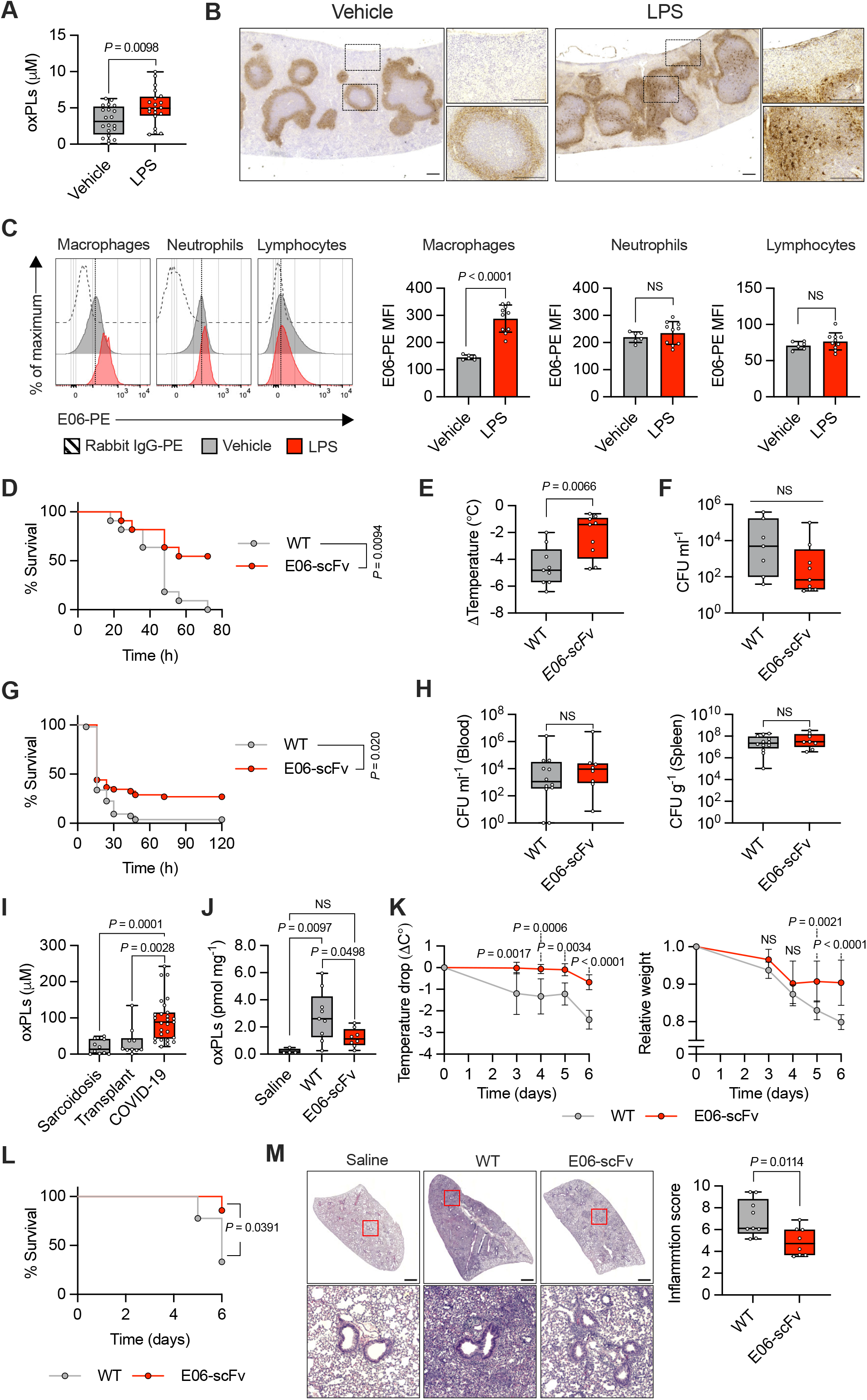
**A**) oxPLs quantification in serum of mice treated with vehicle (saline) or LPS (2 mg Kg^−1^) for 8h. *n* = 20 mice per group. Graph shows means ± SD. Statistical significance was calculated using two-tailed *t* test. **B**) Spleens of mice treated with vehicle (saline) or LPS (2 mg Kg^−1^) for 24h were immunostained with E06 antibody (brown). Representative photomicrographs are shown. Scale bar = 200 μm. **C**) Representative histograms (*left*) and quantification (*right*) of oxPLs accumulation (E06-PE MFI) in the indicated cell populations of the spleens of mice treated with vehicle (saline) or LPS (2 mg Kg^−1^) for 24h. *n* = 6 mice (vehicle), *n* = 10 mice (LPS). Graphs show means ± SD. Statistical significance was calculated using two-tailed *t* test. **D-F**) WT (*n* = 11) and E06-scFv (*n* = 11) mice were subjected to CLP. Survival was followed over time. Kaplan–Meier curves with log-rank (Mantel-Cox) test are shown (**D**). Body temperature loss was measured 8h after surgery (**E**). Bacteria loads in serum were analyzed 24h after CLP (**F**). Graphs show means ± SD. Statistical significance was calculated using two-tailed *t* test. **G-H**) WT (*n* = 53) and E06-scFv (*n* = 52) mice were injected with MRSA (1.8 × 10^8^ CFU/mouse) and monitored for lethality over 120 hours. Kaplan–Meier curves with log-rank (Mantel-Cox) test are shown (**G**). Bacteria loads in serum (*left*) and spleen (*right*) were analyzed 24h after MRSA injection (**H**). Graphs show means ± SD. Statistical significance was calculated using two-tailed *t* test. **I**) oxPLs levels were measured in the BAL of COVID-19 (n = 27) and non-microbially infected (subjects affected by sarcoidosis, n = 10 and transplant, n = 10) patients. Graphs show mean ± SD. Statistical significance was calculated using two-tailed *t* test. **J-M**) WT (*n* = 9) and E06-scFv (*n* = 8) mice were intratracheally administered with poly(I:C) (2.5 mg Kg^−1^) or saline (WT, n = 4) daily for 6 days. oxPLs were quantify from lung homogenates. Graph shows means ± SD. Statistical multiple comparisons were calculated by one-way ANOVA and Tukey’s test (**J**). Body temperature (*left*) and weight (*right*) loss were measured at the indicated time points. Graphs show means ± 95% CL. Statistical comparisons were calculated by two-way ANOVA and Sidak’s test (**K**). Survival was followed over time. Kaplan–Meier curves with log-rank (Mantel-Cox) test are shown (**L**). Representative H&E images of control (saline) and poly(I:C) WT and E06-scFv treated mice. Scale bar = 1 mm. (*left*). Histological inflammation score (*right*). Graph shows means ± SD. Statistical significance was calculated using two-tailed *t* test (**M**).

To test whether oxPLs affected the outcome of the immune response elicited by microbial encounter, we performed cecal ligation and puncture (CLP) to induce a systemic polymicrobial infection in wild-type (WT) mice, or in mice that express the single-chain fragment variable (scFv) of the E06 antibody (E06-scFv mice), which efficiently and specifically neutralizes phosphocholine-containing oxPLs *in vivo* [12, 13, 22]. We initially confirmed that oxPLs were produced and efficiently neutralized in the spleen of E06-scFv, compared to WT, mice subjected to CLP (**Figure S1D**). WT mice in which the activity of oxPLs was not neutralized showed significant increase in mortality and morbidity (as measured by temperature loss), but a similar microbial burden to E06-scFv mice (**Figure 1D-F**).

The CLP model is driven by a polymicrobial infection and takes >2 days to cause death. To further explore the impact of neutralizing oxPLs in an even more inflammatory model, we injected a lethal dose of MRSA sufficient to kill most mice within the first 24h. In keeping with the capacity of oxPLs to increase lethality in the CLP model, mice in which oxPLs were not neutralized showed a significant increased mortality compared to E06-scFv mice when injected with MRSA (**Figure 1G**). When bacterial titers were measured, no significant difference was found in WT, compared to E06-scFv, mice (**Figure 1H**).

Previous studies found increased levels of oxPLs during influenza A virus or severe acute respiratory syndrome coronavirus (SARS-CoV) infections [14, 15]. We, thus, tested whether oxPLs were also augmented in the bronchoalveolar lavages (BALs) of patients infected with SARS-CoV-2, compared to individuals that were transplanted or had sarcoidosis. The samples we utilized were previously characterized for the presence of interferons and inflammatory cytokines, and showed a strong inflammatory response in SARS-CoV-2, but not transplanted or sarcoidosis, patients [23]. We found that the BALs of patients with severe-to-critical coronavirus disease 2019 (COVID-19) contained significantly increased amounts of oxPLs compared to the controls (**Figure 1I**). We, next, utilized a mouse model based on repetitive injections of the synthetic viral ligand polyinosinic-polycytidylic acid (poly(I:C)) to mimic a persistent lung virus infection. This viral mimic was previously utilized to study severe COVID-19 [24] or post-acute syndrome COVID-19 (PASC) [25]. We confirmed that, upon repetitive injections of poly(I:C), the levels of oxPLs in the lungs of WT mice were significantly increased compared to mice that were not injected with poly(I:C), or to E06-scFv mice that received poly(I:C) (**Figure 1J**). Morbidity (measured as temperature or weight loss) and mortality were significantly increased in mice in which the activity of oxPLs was not neutralized (**Figure 1K, L**). Increased mortality was associated with augmented inflammation in the lungs of WT, compared to E06-scFv, mice, as determined by histology (**Figure 1M**).

Overall, our data demonstrate that endogenous oxPLs cause the death of the host upon encounter with pathogens or following chronic stimulation of PRRs. Also, we demonstrate that oxPLs contribute to the immunopathology elicited upon microbial detection independently of the load of the invading microorganisms.

### oxPLs worsen a polymicrobial infection by inhibiting IL-10

Next, we thought to measure global transcriptional changes in macrophages exposed to a microbial ligand and/or to oxPLs to identify inflammatory programs governed by oxPLs that can explain how blocking oxPLs *in vivo* significantly enhances host survival upon microbial encounter. As a source of oxPLs, we utilized the oxidation products of the common phospholipid 1-palmitoyl-2-arachidonyl-sn-glycero-3-phosphocholine (oxPAPC). Bone marrow-derived macrophages (BMDMs) were treated with lipopolysaccharide (LPS) for 3 hours, followed by oxPAPC treatment for 3 or 18 hours, and Illumina HiSeq RNA sequencing (RNAseq) was performed (**Figure 2A**). In keeping with our previous findings [20], *Il1b* was upregulated late upon oxPAPC administration in the presence of LPS. However, we also found a set of genes that were downregulated early, and remained suppressed also at late time points. Among these genes, we focused on the key anti-inflammatory cytokine IL-10 [26] and confirmed that oxPAPC inhibits the transcription of *Il10* in macrophages activated with LPS (**Figure 2B**).

**Figure 2:**
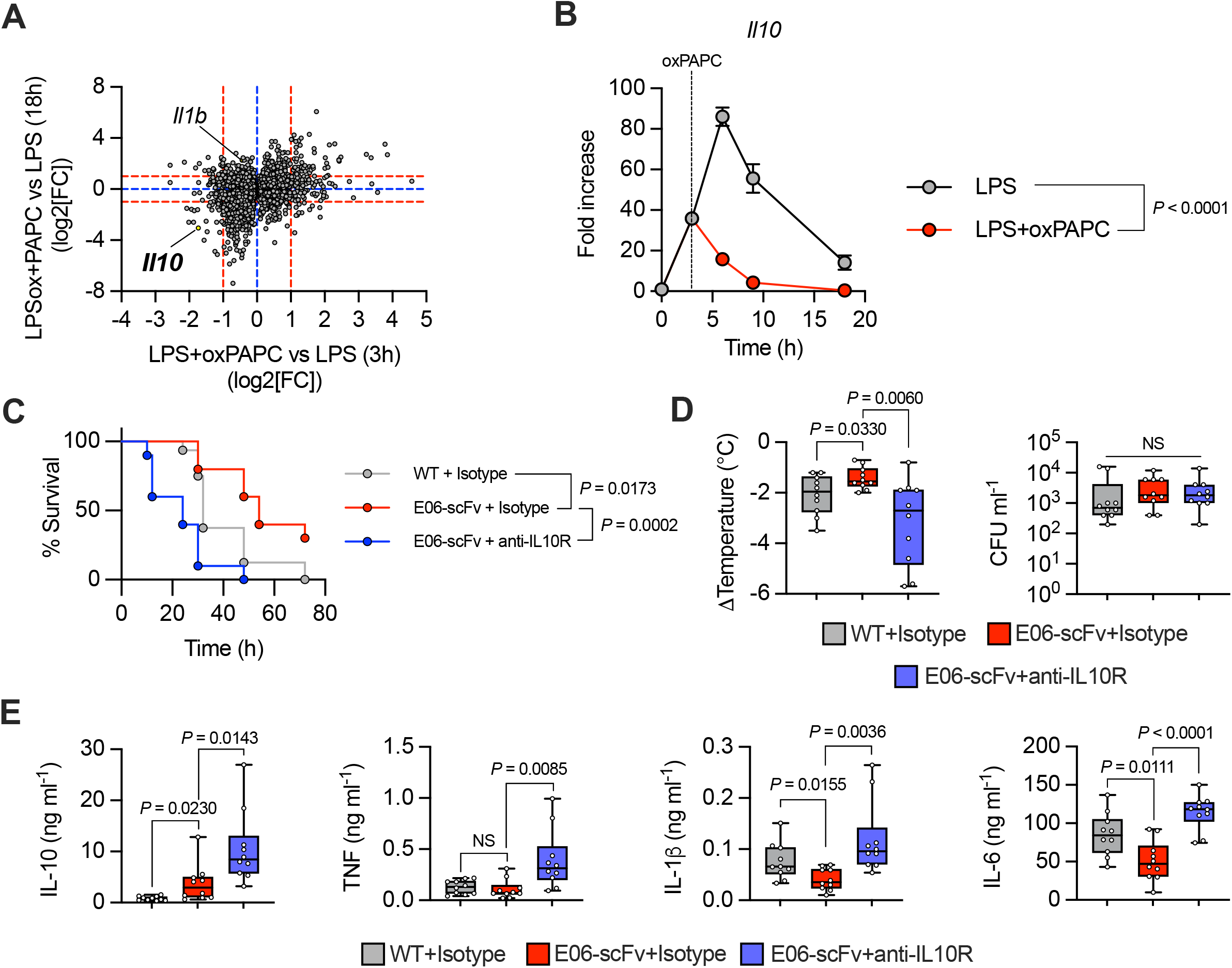
**A**) Scatter plot of transcript fold-change expression with s-value < 0.01 (RNA-seq) in LPS-primed BMDMs treated, or not, with oxPAPC (100 μg ml^−1^) for 3h (x-axis) or 18h (y-axis). Points correspond to genes. **B**) BMDMs were primed with LPS (1 μg ml^−1^) and then treated, or not, with oxPAPC (100 μg ml^−1^). *Il10* mRNA levels were measured at the indicated time points by qPCR. *n* = 3, statistical significance was calculated using two-way ANOVA and Sidak’s multiple comparisons test. **C-E**) WT (*n* =16) and E06-scFv mice (*n* =10, isotype control *n* = 10, anti-IL10R) undergoing CLP were treated with anti-IL-10R antibody or isotype control, as indicated. Survival was followed over time. Kaplan–Meier curves with log-rank (Mantel-Cox) test are shown (**C**). (**D**) Body temperature loss was measured 8h after surgery (left) and bacteria loads in serum were analyzed 24h after CLP (right). Statistical significance was calculated using two-tailed *t* test. (**E**) WT and E06-scFv mice undergoing CLP were treated with anti-IL-10R antibody or isotype control, as indicated, and serum levels of IL-10, TNF, IL-1β and IL-6 were analyzed 8h after CLP. *n* =10 mice per group. Graph show means ± SD. Statistical significance was calculated using two-tailed *t* test.

To account for the reduced pathology and increased survival in in E06-scFv, compared to WT, mice in our three models of microbial-driven inflammation (**Figure 1D-M**), we hypothesized that the inhibition of the activity of oxPLs in E06-scFv mice increased IL-10 levels. Indeed, we found that levels of IL-10 were significantly increased in E06-scFv, compared to WT, mice in all the three models utilized (**Figure S2A-C**). In contrast, levels of pro-inflammatory cytokines were negatively regulated, or left unaltered, in E06-scFv, compared to WT, mice (**Figure S2A-C**).

These data support the hypothesis that oxPLs increase mortality upon microbial encounter by inhibiting IL-10 production. To test whether inhibition of IL-10 by oxPLs is the cause of the death of mice, we inhibited IL-10 signaling in E06-scFv mice. For this experiment, we focused on the CLP polymicrobial infection model that allows to follow the mice over time and to detect not only pro- and anti-inflammatory cytokines, but also to test pathogen control by assessing microbial titers. We found that blockade of IL-10 signaling in E06-scFv mice reverted their phenotype, increasing morbidity, mortality, and restoring levels of pro-inflammatory cytokines similar to WT mice, without affecting bacterial count (**Figure 2C-E**).

Overall, these data demonstrate that naturally occurring oxPLs produced in response to microbial encounters exacerbates immunopathology by reducing the levels of IL-10.

### oxPLs regulate IL-10 levels by inhibiting AKT

To determine the molecular mechanisms that leads to IL-10 inhibition upon oxPLs encounter, we utilized oxPAPC on BMDMs. Since we previously found that oxPAPC alters the metabolism of macrophages [20], we tested whether the rapid change in the transcription of *Il10* that follows oxPAPC administration is driven by changes in the respiratory and/or glycolytic capacity of macrophages. We found that oxPAPC, but not dipalmitoylphosphocholine (DPPC) (a phospholipid that cannot be oxidized and is immunologically inactive [18–20]), rapidly inhibited glycolysis, in the presence or absence of LPS (**Figure 3A, S3A**), while OXPHOS was only marginally affected (**Figure 3B, S3B**). To test whether glycolysis inhibition drives the reduction of IL-10, we inhibited the activity of hesokinase-2 or of lactate dehydrogenase in LPS-treated cells exposed (or not) to oxPAPC. We found that inhibition of the glycolytic cascade did not cause a change in the levels of IL-10 (or TNF, used as a control cytokine, which levels are not affected by oxPAPC [27]) either in the presence or absence of oxPAPC stimulation (**Figure S3C, D**). In keeping with these results, culturing macrophages in a low glucose medium did not affect the levels of IL-10 in response to LPS and/or oxPAPC stimulation (**Figure S3E**).

**Figure 3:**
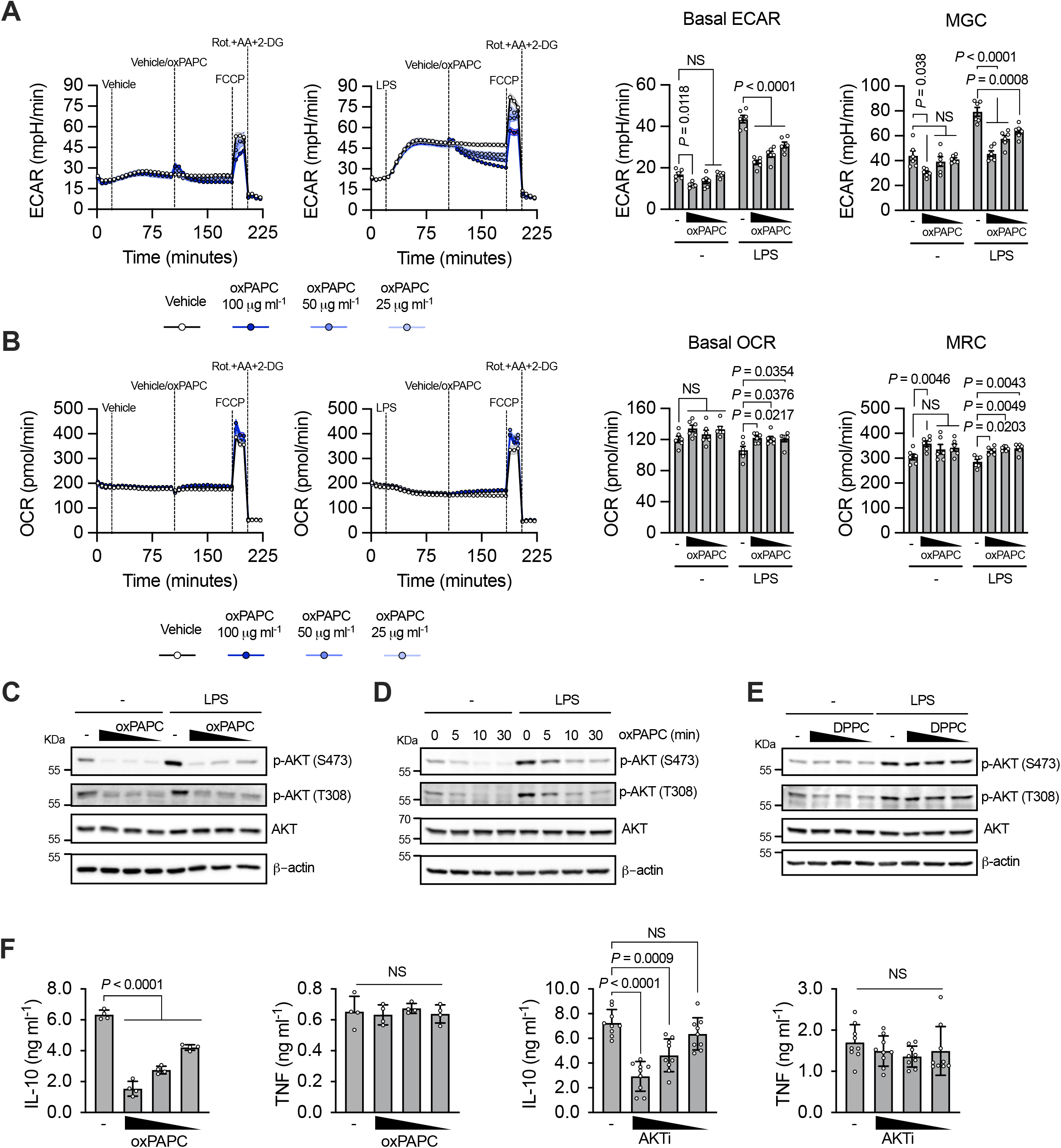
**A**) Real-time changes in ECAR (*left*), basal ECAR and maximal glycolytic capacity (MGC) quantifications (*right*) of BMDMs untreated or treated with LPS (1 μg ml^−1^) and then challenged with different doses of oxPAPC, as indicated. **B**) Real-time changes in OCR (*left*), basal OCR and maximal respiratory capacity (MRC) quantifications (*right*) of BMDMs treated as in **A**. *n* = 6, graphs are representative of three independent experiments and show means ± SEM. Statistical significance was calculated using two-way ANOVA and Dunnett’s multiple comparisons test. **C**) BMDMs were primed, or not, with LPS and then stimulated with oxPAPC (100, 50 or 25 μg ml^−1^). AKT phosphorylation was analyzed after 1h by immunoblotting. **D**) BMDMs were primed, or not, with LPS and then stimulated with oxPAPC (100 μg ml^−1^). AKT phosphorylation was analyzed at the indicate time points by immunoblotting. **E**) BMDMs were primed, or not, with LPS and then treated with DPPC (100, 50 or 25 μM). AKT phosphorylation was analyzed after 1h by immunoblotting. Images are representative of three independent experiments. **F**) BMDMs were primed with LPS and then stimulated with oxPAPC (100, 50 or 25 μg ml^−1^) (*n* = 4) (*left*) or AKTi (10, 5, 25 μM) (*n* = 9) (*right*). IL-10 and TNF production was quantified by ELISA 18h later. Graphs are representative of four independent experiments and show means ± SEM. Statistical multiple comparisons were calculated by two-way ANOVA and Dunnett’s test.

We next tested whether oxPAPC alters the activation of AKT, which is the first enzyme that controls the early wave of glycolysis downstream of TLR4 [28, 29], but it is also a master regulator of multiple inflammatory processes [30]. oxPAPC, but not DPPC, administration, in the presence or absence of LPS, inhibited AKT phosphorylation in a dose- and time-dependent manner (**Figure 3C-E**). Targets and downstream kinases of AKT were also inhibited by oxPAPC, confirming that oxPAPC inhibits the activity of AKT (**Figure S3F**). Of note, administration of the AKT inhibitor VIII (AKTi) mirrored the capacity of oxPAPC to inhibit IL-10 (but not TNF) production and glycolysis (**Figure 3F, S3G, H**).

We next assessed whether oxPAPC maintained its capacity to inhibit AKT and dampen IL-10 production in phagocytes of different origin. Mouse dendritic cells (DCs) were differentiated *in vitro* using GM-CSF. Alternatively, primary mouse macrophages were isolated from the peritoneum. The capacity of oxPAPC to inhibit AKT, glycolysis, and IL-10, but not TNF, production was assessed. Our data demonstrate that oxPAPC maintains its activity on the different mouse phagocytes (**Figure S3I-N**).

Finally, we tested the activity of oxPAPC when cells were activated with different PAMPs. We utilized PAMPs of bacterial, viral, and fungal origin and found that, regardless of the PRR stimulated, oxPAPC maintained its inhibitory activities (**Figure S4A-F**). Of note, in macrophages stimulated with type I interferons, oxPAPC inhibited AKT and the induction of *Il10* (while other interferon-stimulated genes were only marginally affected) (**Figure S4G, H**). AKT inhibition also dampened the transcription of *Il10* in cells stimulated with type I interferons (**Figure S4I**).

Overall, these data demonstrate that oxPAPC leads to AKT inhibition, and consequently blocks IL-10 production, in response to distinct PAMPs and in multiple mouse phagocytes.

### oxPLs exert their activities in human phagocytes and are inversely correlated with IL-10 levels in patients with polymicrobial infections

We next tested whether oxPAPC also acts on human phagocytes. Human monocytes, monocyte-derived-DCs or monocyte-derived macrophages were stimulated with LPS in the presence or absence of oxPAPC. For all human cells, oxPAPC maintained the capacity to dampen AKT activation, glycolysis induction, and the production of IL-10, but not of TNF (**Figure 4A-I**).

**Figure 4.**
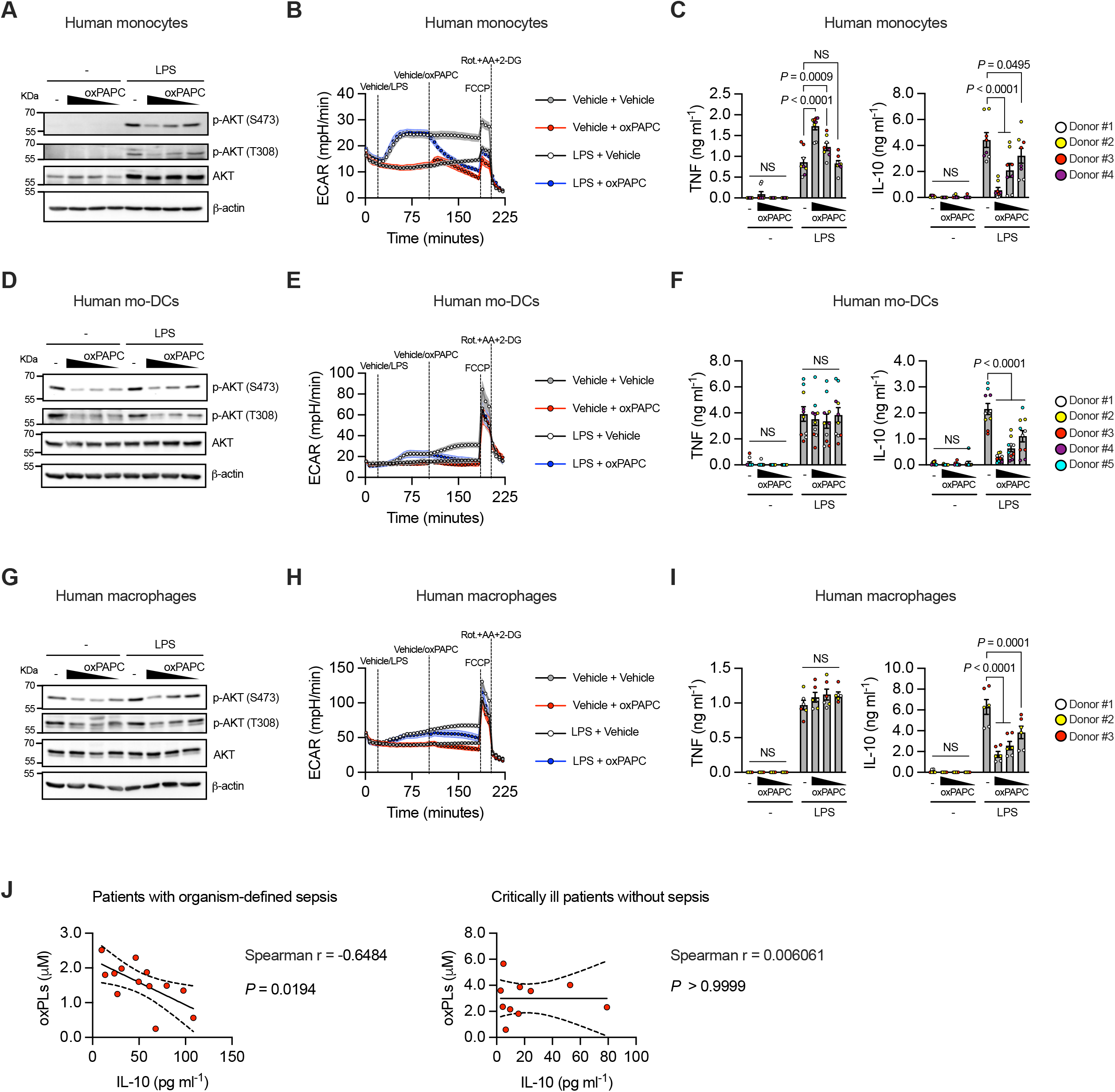
**A**) Human monocytes were primed, or not, with LPS and treated or not with oxPAPC (50, 12 or 12.5 μg ml^−1^) for 1h. AKT phosphorylation was assessed by immunoblot. Data are representative of three independent experiments. **B**) ECAR of human monocytes, treated as indicated, was measured using a Seahorse analyzer. oxPAPC was used at 50 μg ml^−1^. Data are representative of three independent experiments. **C**) Human monocytes were treated as in **A** and TNF and IL-10 release was quantified by ELISA after 18h. *n* = 8, from 4 different donors. Graphs show means ± SEM. Statistical multiple comparisons were calculated by two-way ANOVA and Dunnett’s test. **D**) Human mo-DCs were primed, or not, with LPS and treated or not with oxPAPC (50, 12 or 12.5 μg ml^−1^) for 1h. AKT phosphorylation was assessed by immunoblot. Data are representative of three independent experiments. **E**) ECAR of human mo-DCs, treated as indicated, was measured using a Seahorse analyzer. oxPAPC was used at 50 μg ml^−1^. Data are representative of three independent experiments. **F**) Human mo-DCs were treated as in **D** and TNF and IL-10 release was quantified by ELISA after 18h. *n* = 10, from 5 different donors. Graphs show means ± SEM. Statistical multiple comparisons were calculated by two-way ANOVA and Dunnett’s test. **G**) Human macrophages were primed, or not, with LPS and treated or not with oxPAPC (50, 12 or 12.5 μg ml^−1^) for 1h. AKT phosphorylation was assessed by immunoblot. Data are representative of three independent experiments. **H**) ECAR of human macrophages was measured using a Seahorse analyzer. oxPAPC was used at 50 μg ml^−1^. Data are representative of three independent experiments. **I**) Human macrophages were treated as in **G** and TNF and IL-10 release was quantified by ELISA after 18h. *n* = 6, from 3 different donors. Graphs show means ± SEM. Statistical multiple comparisons were calculated by two-way ANOVA and Dunnett’s test. **J**) Scatter plot of oxPLs and IL-10 levels in plasma of patients with organism-defined sepsis (*left*, n = 13) and critically ill patients without sepsis (*right*, n = 14), obtained within 48 hours of admission to the intensive care unit. Plots show results of Spearman’s rank correlation tests and a linear regression line with 95% confidence intervals.

Based on these data, we investigated whether the presence of oxPLs in subjects with polymicrobial infections was correlated with the levels of IL-10. We focused on pediatric individuals sampled when they entered the intensive care unit (ICU) for evaluation of possible sepsis. Hypotension requiring vasopressor medications and identification of infectious pathogens occurred more frequently in patients meeting diagnostic criteria for sepsis [31] compared to those without sepsis. Patient demographics, respiratory compromise, ventilatory support, and glucocorticoid use were comparable between the two groups. Of note, oxPAPC levels were inversely correlated with circulating IL-10 levels only in the cohort of critically infected patients with a confirmed diagnosis of sepsis (**Figure 4J**).

Overall, these data demonstrate that the capacity of oxPLs to block AKT activation and glycolysis, and to inhibit IL-10 production is maintained across multiple human phagocytes. Our data also show that critical patients with polymicrobial infections present reduced levels of IL-10 when the blood concentration of oxPLs is increased.

### oxPLs directly bind and inhibit AKT

We next determined how oxPAPC inhibits AKT. We initially tested whether oxPAPC administration affects the three major pathways that activate AKT, namely PI3K, TBK1/IKKε, and PDK1 [28, 30] (**Figure S5A**). We found that none of these pathways was altered by oxPAPC administration to macrophages treated, or not, with LPS (**Figure S5B**). Of note, we did not reveal any major change in the activity of the complex 1 or 2 of mTOR (**Figure S5C**). We, thus, tested whether any of the known receptors of oxPAPC [5, 17, 32–39] were involved in the inhibition of AKT, glycolysis, and IL-10 induction. We excluded the involvement of TLR4 and TLR2 (**Figure 5A, B**), and of CD14 and CD36 (**Figure 5C, D**). Although we confirmed that oxPAPC activates NRF2 in macrophages [38] (**Figure S5D**), we found that oxPAPC maintained the capacity to inhibit AKT, glycolysis, and IL-10 in *Nfe2l2*^−/−^ cells (**Figure 5E**). In keeping with the fact that NRF2 is not involved in regulating AKT and IL-10, administration of the NRF2 activator 15-Deoxy-Δ^12,14^-prostaglandin J2 (15d-PGJ_2_) [40] did not affect AKT activation or IL-10 levels in LPS-stimulated macrophages (**Figure S5E, F**).

**Figure 5:**
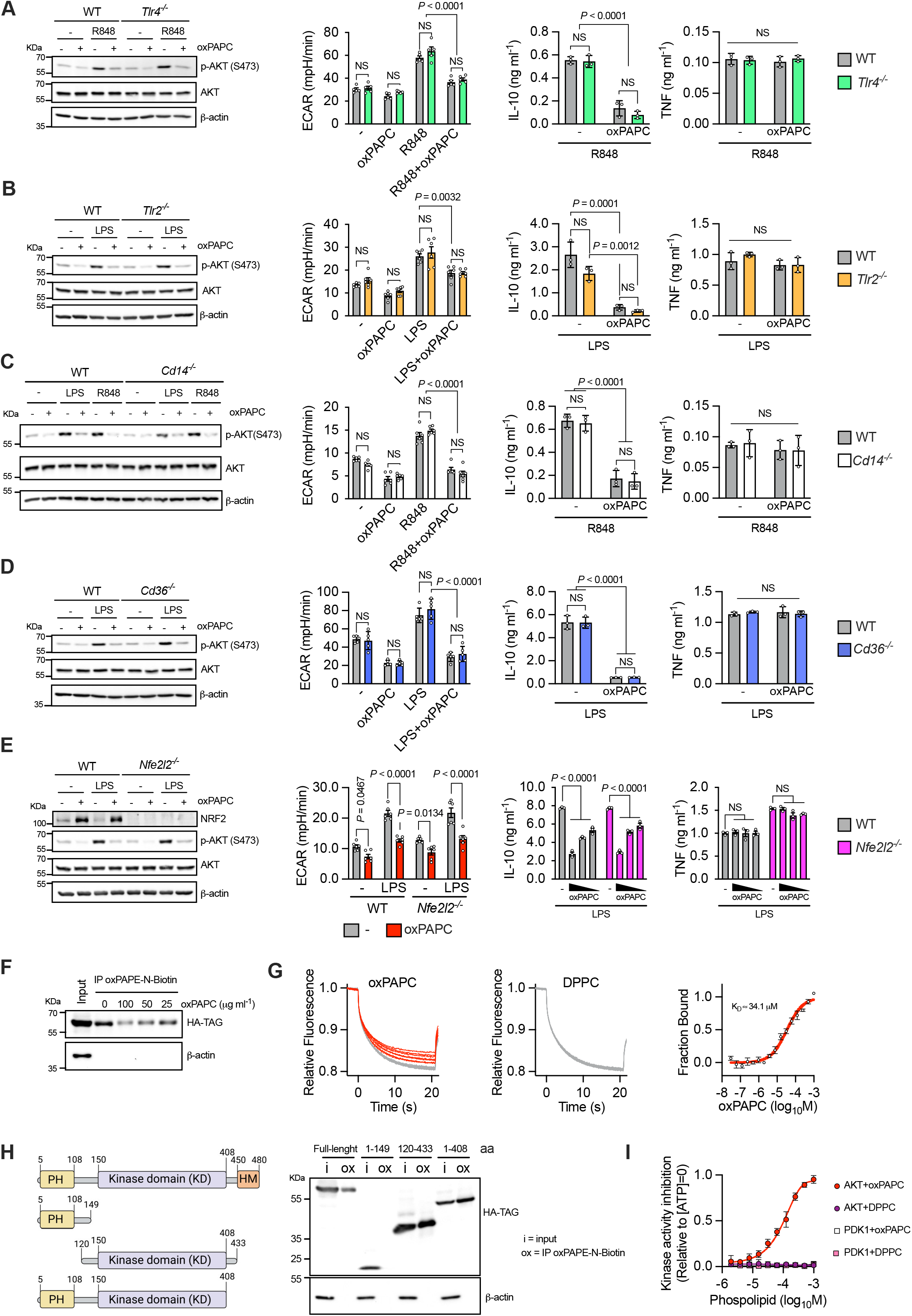
**A-E**) WT and *Tlr4^−/−^*(**A**), *Tlr2^−/−^* (**B**), *Cd14^−/−^* (**C**), *Cd36^−/−^* (**D**) *or Nfe2l2^−/−^* (**E**) BMDMs were primed, or not, with R848 (1 μg ml^−1^) (**A, C**) or LPS (1 μg ml^−1^) (**B-E**) and treated or not with oxPAPC (100 μg ml^−1^) for 1h (*left, center*) or 24h (*right*). AKT phosphorylation (**A-E**) or NRF2 accumulation (**E**) were assessed by immunoblot (*left*). Data are representative of three independent experiments. ECAR (*center*) was measured using a Seahorse analyzer. *n* = 6 (**A-C, E**), n = 5 (**D**), data are representative of three independent experiments and show means ± SEM. Statistical multiple comparisons were calculated by two-way ANOVA and Tukey’s test. IL-10 and TNF release (*right*) was quantified by ELISA. *n* = 3, graphs are representative of three independent experiments and show means ± SEM. Statistical multiple comparisons were calculated by two-way ANOVA and Tukey’s test. **F**) oxPAPC binding capacity of AKT was determined by pull down assay. Cellular lysate of 293T cells expressing HA-tagged human AKT1 was incubated with oxPAPE-N-Biotin and the indicated doses of oxPAPC. AKT associated with biotinylated lipids was captured by streptavidin beads and revealed by immunoblotting using anti-HA antibody. β-actin was used as a negative control. Data are representative of three independent experiments. **G**) Microscale thermophoresis (MST) analysis of oxPAPC and AKT interactions. Traces of fluorescently labeled human recombinant AKT1 incubated with oxPAPC (1000, 500, 250,125, 62.5, 31.25, 15.62, 7.8, 3.9, 1.9, 0.97, 0.488, 0.24, 0.12, 0.0061 and 0.030 μM) or DPPC (500, 250,125, 62.5, 31.25, 15.62, 7.8, 3.9, 1.9, 0.97, 0.488, 0.24, 0.12, 0.0061 and 0.030 μM) (*left and central*). oxPAPC-AKT binding curve was derived from the quantification of normalized fluorescence changes (*right*). *n* = 3, graph shows means ± SD. Images are representative of three independent experiments. **H**) Cellular lysate of 293T cells expressing the indicated HA-tagged AKT1 truncated forms were incubated with oxPAPE-N-Biotin. HA-proteins associated with biotinylated lipid were captured by streptavidin beads and revealed by immunoblotting. Data shown are representative of three independent experiments. **I**) Active human recombinant AKT1 or PDK1 were incubated with oxPAPC or DPPC (1000, 500, 250,125, 62.5, 31.25, 15.62, 7.8, 3.9, and 1.9 μM) and kinase-specific FRET-peptide substrates (Z-’LYTE). Kinase inhibition was measured as the ability of lipid to block substrate phosphorylation. *n* = 4, graph shows means ± SD. Data are representative of three independent experiments.

Based on these data, we tested the hypothesis that oxPAPC can directly bind and inhibit AKT. Initially, we utilized a biotinylated analog of oxPAPC (oxPAPE-N-biotin) to pull down a HA-tagged AKT. The experiment was performed in the presence or absence of oxPAPC to compete for the binding to AKT. Our data demonstrated that oxPAPC and AKT interact with each other (**Figure 5F**). Next, to assess whether oxPAPC directly binds AKT, we used MicroScale thermophoresis (MST) [41]. Based on the Immgen database, tissue resident macrophages mostly express *Akt1* and *Akt2*, except for peritoneal macrophages that also express *Akt3* (**Figure S5G**). We confirmed that bone-marrow derived macrophages express all three isoforms of AKT (**Figure S5H**). We, thus, tested the capacity of oxPAPC to bind each of the three AKT isoforms and we found that oxPAPC, but not DPPC, directly binds AKT1-3 with a K_d_ that varies between 22.7μM and 65.3μM, depending on the AKT isoform (**Figure 5G, S5I, J**). To identify the domain of AKT bound by oxPAPC, we expressed the following HA-tagged AKT1 proteins: i) full length; ii) amino acid (AA)1-149 (PH domain); iii) AA120-433 (KD domain); and iv) AA1-408 (PH and KD domains) (**Figure 5H**). These proteins were then tested for their ability to interact with oxPAPE-N-biotin. We found that oxPAPC binds the catalytic domain of AKT (**Figure 5H**).

Since oxPAPC binds to the catalytic domain of AKT, we assessed whether oxPAPC directly inhibits AKT activity. The enzymatic activity of AKT1, 2 and 3 was tested in the presence of oxPAPC (or DPPC) in a cell-free assay. oxPAPC, but not DPPC, inhibited the activity of AKT1-3, but not of the related kinase PDK1, in a dose-dependent manner (**Figure 5I, S5K, L**).

Overall, these data demonstrate that oxPAPC directly binds and inhibits all three isoforms of AKT.

### oxPLs-dependent AKT inhibition reprograms the landscape of metabolites in macrophages

We next investigated how oxPAPC-dependent AKT inhibition affects IL-10 production. Cells were stimulated with LPS, and the main transcriptional pathways activated downstream of the Myddosome were assessed (**Figure S6A**). We found that canonical NF-kB activation was not altered in LPS-stimulated cells exposed to oxPAPC (**Figure S6B, C**). Similarly, the ERK, JNK, and p38 pathways were not altered by oxPAPC (**Figure S6D**). Previous reports implicated CREB in AKT-dependent regulation of IL-10 [42, 43], but we did not detect changes in CREB activation in the presence of oxPAPC in LPS-stimulated cells (**Figure S6D**). Similar to Myddosome dependent signaling, oxPAPC did not alter TRIF- and IRF3-dependent responses (**Figure S6E-G**). These results are in agreement with our previous data showing that oxPAPC activity happens across multiple stimuli and independently of distinct PRRs.

To identify the pathway that regulates IL-10 production upon oxPAPC-dependent inhibition of AKT, we interrogated the landscape of metabolites in cells activated, or not, with LPS, and exposed, or not, to oxPAPC, or to AKTi. A sparse partial least squares discriminant analysis (sPLS-DA) revealed that the various treatments induced a unique landscape of metabolites (**Figure 6A, B**). Also, a pathway analysis revealed the regulation of several metabolic pathways, some of which were shared by cells exposed to oxPAPC or AKTi (**Figure 6C**). Several of the common metabolic pathways, such as “Purine” or “Serine/glycine” metabolism, converge on the methionine cycle (**Figure 6D**). S-adenosyl-homocysteine (SAH) and/or homocysteine, which are intermediates of the methionine cycle, were among the metabolites most upregulated in cells exposed to LPS and oxPAPC, or AKTi (**Figure 6E**). Notably, the conversion of S-adenosyl-methionine (SAM) into SAH/homocysteine liberates methyl groups that can be used for epigenetic modifications such as H3K27me2me3, a central modulator of gene silencing in mammals [44, 45]. Given that AKT suppresses the activity of the methyltransferase EZH2 (which methylates H3K27 [46]), we hypothesized that oxPAPC-dependent AKT inhibition enhances EZH2 activity, and thereby suppresses *Il10* transcription.

**Figure 6.**
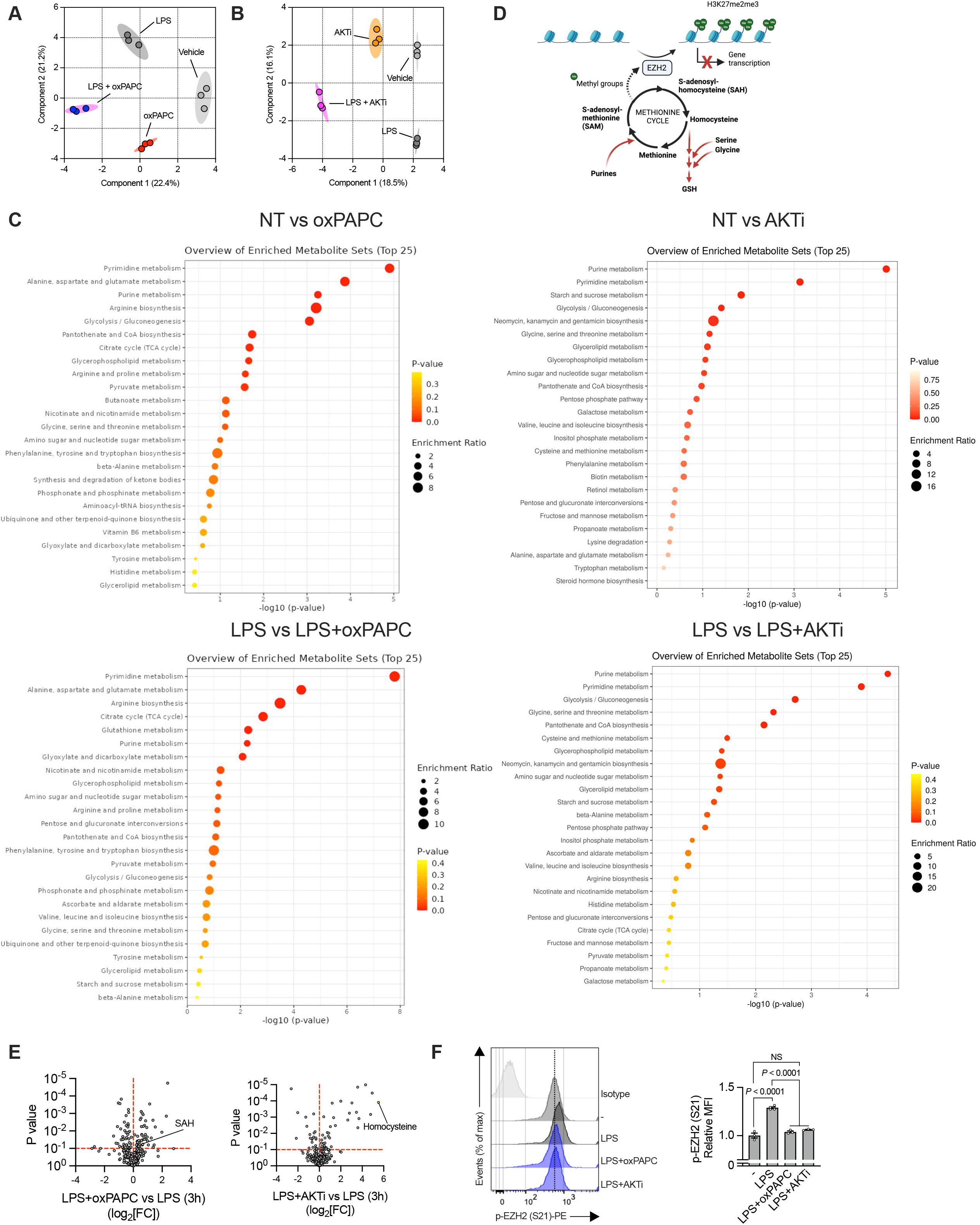
**A-C**) BMDMs were primed, or not, with LPS (1 μg ml^−1^) treated or not with oxPAPC (100 μg ml^−1^) or AKTi (10 μM). Mass-spectrometry polar metabolomics analysis was performed 3h later. sPLS-DA plot showing the effect on variance during oxPAPC (**A**) or AKTi (**B**) treatments. Each point represents an independent biological sample. KEGG pathway enrichment analysis of the indicated treatments (**C**). **D**) Schematic showing the interconnection between methionine cycle and histone methylation processes. **E**) BMDMs were primed with LPS and then stimulated with oxPAPC (100 μg ml^−1^) (*n* = 3) (left) or AKTi (10 μM) (*n* = 3) (right). Volcano plots show the change of polar metabolites after 3h. **F**) BMDMs were treated as in **E** and EZH2 phosphorylation was measured by flow cytometry after 1h. *Left*: cytofluorimetry histograms. *Right*: bar graphs. Bars represent the MFI of p-EZH2. *n* = 3, graph shows means ± SEM. Statistical multiple comparisons were calculated by two-way ANOVA and Tukey’s test. Data are representative of three independent experiments.

oxPAPC administration, as well as AKTi treatment, reduced the inhibitory phosphorylation of EZH2 on S21 [46] (**Figure 6F**). Furthermore, our RNAseq analysis showed that *Id3*, a gene activated by EZH2 [47], was upregulated in oxPAPC-treated cells.

Overall, our data demonstrate that oxPAPC rewires the metabolic landscape of LPS-stimulated macrophages, suggesting that epigenetic changes can be the cause of the inhibition in the induction of *Il10*.

### oxPLs-dependent AKT inhibition epigenetically reprograms Il10 transcription

To directly test the capacity of oxPAPC to alter H3K27me2me3 in the *Il10* locus, we performed chromatin immune precipitation (ChIP) followed by qPCR analysis of several conserved non-coding sequences (CNS) (**Figure 7A**). H3K27me2me3 was highly enriched in *Hoxc10* and poorly present in *Gadph*, utilized as positive and negative controls, respectively, independently of cell stimulation. We found that oxPAPC treatment significantly restored the levels of H3K27me2me3 in multiple CNS of cells treated with LPS and exposed to oxPAPC, compared to cells treated with LPS alone (**Figure 7A**). Of note, inhibition of the activity of EZH2 with two different highly selective methyltransferase inhibitors [48, 49], as well as with an EED degrader [50], restored IL-10 production in cells treated with LPS and oxPAPC, only minimally impacting TNF levels (**Figure 7B, S7A-C**). In keeping with the *in vitro* activity of oxPAPC on EZH2, when EZH2 was inhibited *in vivo*, mice were protected against the polymicrobial infection that follows CLP (**Figure 7C, D**) and IL-10 levels were increased, while, as a consequence of IL-10 activity, pro-inflammatory cytokines levels were dampened (**Figure 7E**). As shown before, bacterial levels were not significantly affected (**Figure 7F**), demonstrating that targeting oxPL-dependent activation of EZH2 ameliorates disease development, without altering the anti-microbial activity of the immune system.

**Figure 7:**
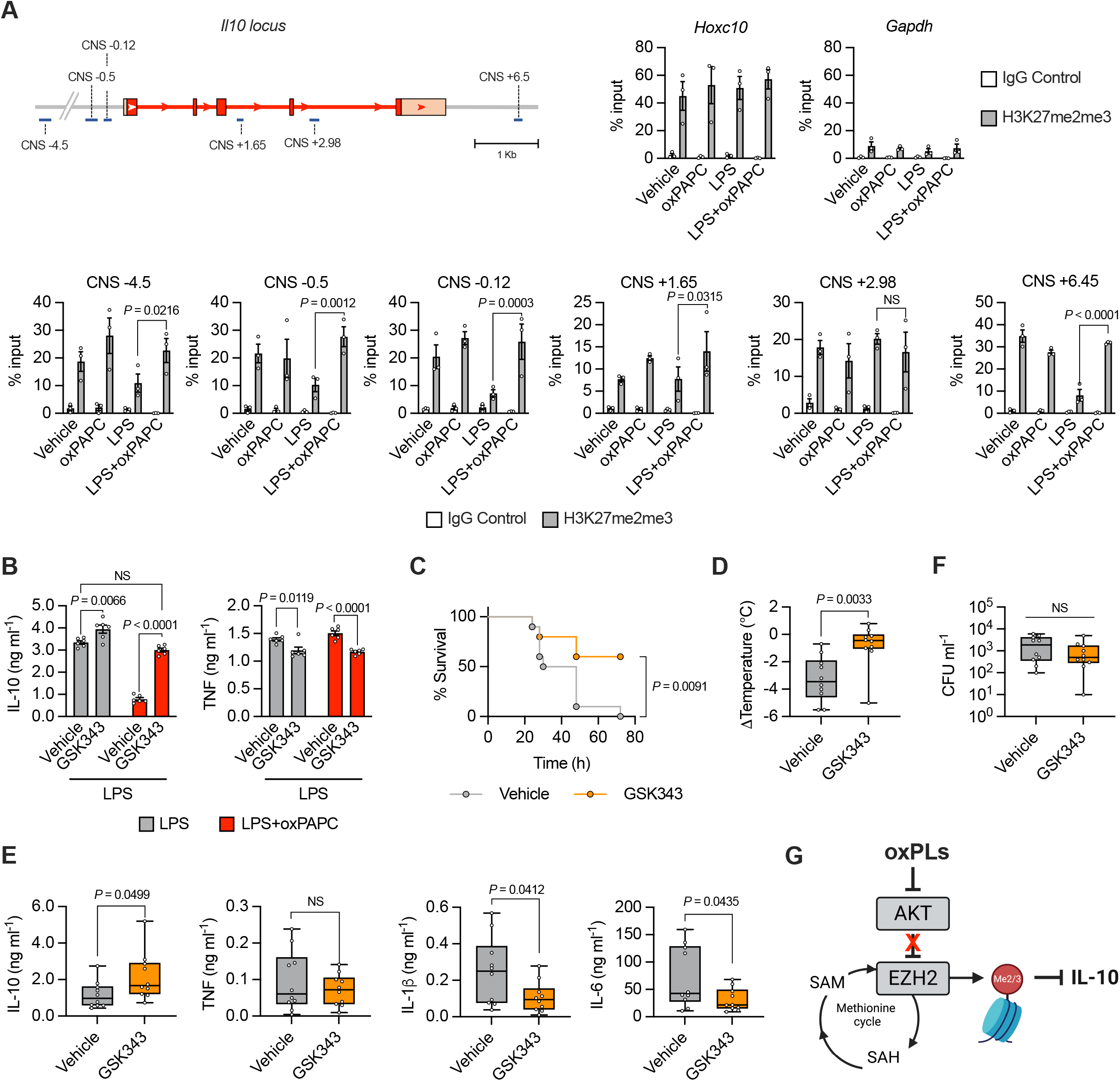
**A**) BMDMs were primed, or not, with LPS and then stimulated with oxPAPC (100 μg ml^−1^). Chromatin immunoprecipitation and quantitative PCR (ChIP–qPCR) of H3K27me2me3 at CNS regions related to the *Il10* locus (indicated in the scheme), was performed 3h later. *Hoxc10* and *Gapdh* were used as positive and negative controls, respectively. *n* = 3 independent immunoprecipitations, graphs show means ± SEM. Statistical significance was calculated using two-way ANOVA. **B**) BMDMs were incubated with GSK343 (10 μM) for 30 minutes, primed with LPS and then stimulated with oxPAPC (100 μg ml^−1^). IL-10 and TNF production was quantified by ELISA after 18h. *n* = 6, graphs are representative of four independent experiments and show means ± SEM. Statistical multiple comparisons were calculated by two-way ANOVA and Tukey’s test. **C-F**) WT mice (*n* = 10) were treated with GSK343 and subjected to CLP. Survival was followed over time. Kaplan–Meier curves with log-rank (Mantel-Cox) test are shown (**C**). Body temperature loss was measured 8h later after surgery (**D**). Serum levels of IL-10, TNF, IL-1β and IL-6 were analyzed 8h after CLP (**E**). Bacteria loads in serum were analyzed 24h after CLP (**F**). Statistical significance was calculated using two-tailed *t* test. **G**) Schematic showing how oxPLs control IL-10 production by guarding the activity of AKT.

Overall, our data demonstrate that via the inhibition of AKT, oxPAPC rewires the methionine cycle and boosts the activity of EZH2, which in turns increases the trimethylation of H3K27, thus dampening IL-10 production (**Figure 7G**). Also, that targeting EZH2 *in vivo* counterbalances the activity of oxPLs, ameliorating the immunopathology driven by excessive inflammation that follows microorganism encounter.

## Discussion

oxPLs are host-derived inflammatory cues that are generated in the context of sterile inflammation and inflammatory diseases [16]. In our work, we demonstrate that host-derived oxPLs are formed as a consequence of the encounter with distinct PAMPs, and that oxPLs are key determinants of the outcome of the immune response. In particular, we found that oxPLs reduce the production of IL-10, thus exaggerating the inflammatory response that is secondary to a disseminated polymicrobial infection, to MRSA encounter, and to persistent sensing of a viral mimic in the lung. oxPLs exert this function by inhibiting AKT and rewiring the methionine cycle, which epigenetically switches off the *Il10* locus via the activity of the epigenetic writer EZH2.

By using the E06-scFv transgenic mouse model, whose secreted sc-Fv targets the phosphocholine moieties of oxPLs, we show that oxPLs blockade significantly increased the survival of mice in multiple mouse models of microbial sensing. Our data demonstrate that oxPLs do not compromise immune resistance and, instead, are fundamental to determine the extent of the inflammation elicited in response to microbial sensing. In addition, our findings that targeting oxPLs in distinct infectious models was effective in preventing mortality open to the development of a novel approach against microbial infections, switching the focus of therapeutic intervention from the pathogen to the host. This is particularly relevant in light of our findings that septic patients present an inverse correlation between the levels of oxPLs and circulating IL-10. Under these circumstances, the use of drugs that target the metabolic and epigenetic adaptions initiated by oxPLs during sepsis may become a new valuable therapeutic tool.

From a physiological point of view, our data reveal that oxPLs serve as sensors of tissue damage and alert the immune system to potentiate its response against invading microorganisms. The primary function of the immune system is to prevent pathogen invasion and spread. It is, thus, necessary that the cells of the immune system recognize the presence of host damage caused by the growth of pathogens that are not properly controlled. In this context, the immune system reacts by increasing the potency and extent of the immune response. The “collateral” damage to the tissues of the host caused by the potentiated immune response would be evolutionary justified by the necessity to control the spread of the pathogen. In our *in vivo* experimental models, we mimicked inflammatory conditions that are extreme, as represented by systemic polymicrobial or Gram-positive-driven infections, or a persistent stimulation of the immune response by a viral mimic, similarly to what has been described in the context of severe COVID-19 or PASC [24, 25]. Under these conditions, the excessive inflammation driven by oxPL-dependent inhibition of IL-10 causes the death of the host. In contrast, under less potent inflammatory conditions, the production of oxPLs may tip the balance of the immune response in a way that will help to alert the immune system to eradicate the invading microorganisms, without causing excessive inflammation and lethality.

Our data also showed that none of the known receptors for oxPLs are involved in regulating IL-10 levels. In contrast, the reduction in the production of IL-10 is driven by the block of the “physiological” activity of AKT and by the consequent activation of the epigenetic writer EZH2. Under our experimental conditions, oxPAPC acts as a “*self*” virulence factor that initiates ETI by directly binding and inhibiting AKT. AKT is a key inflammatory and metabolic checkpoint that regulates the immune response [30]. Thus, it is important for the host to “guard” the activity of AKT, and, in case of alteration of its activation, to instruct the immune system to respond adequately. In the case of oxPAPC-dependent inhibition of AKT, the phagocytes of the host respond by decreasing IL-10 production. The reduced activity of IL-10 *in vivo* on bystander cells in turns leads to increased pro-inflammatory cytokine induction. Intriguingly, AKT is the target of multiple virus and bacterial effectors that can either inhibit or boost its activity. For example, AKT activation is blocked by *Yersinia* YopJ, *Pseudomonas* ExoT, *Vibrio cholerae* MakA, and uropathogenic *Escherichia coli* alpha-hemolysin [51–53]. In these contexts, it will be important in future studies to assess whether the host elicits a response similar to the one initiated in response to oxPLs to reduce IL-10. In contrast to virulence factors that block AKT, several bacterial or viral pathogens such as *Salmonella*, *Shigella*, influenza viruses, or herpesviruses exploit the activation of the AKT pathway to prevent cell death and favor proliferation [51, 54, 55]. Under these circumstances, oxPL production may exert anti-microbial roles, and may be evolutionary advantageous. Therefore, it will be important to contextualize the activity of oxPLs *in vivo* based on the pathogen encountered and according to the extent of inflammation elicited.

Overall, our data revealed that the activity of oxPLs can elicit an ETI response, expanding our understanding of the functioning of the immune system and of the contribution of external- and self-derived inflammatory cues to controlling the extent of the inflammatory response.

## Acknowledgements

IZ is supported by NIH grants 2R01AI121066, 2R01DK115217, 1R01AI165505, 1R01AI170632, and contract no. 75N93019C00044, Lloyd J. Old STAR Program CRI3888, and holds an Investigators in the Pathogenesis of Infectious Disease Award from the Burroughs Wellcome Fund. MDG is supported by The Crohn’s & Colitis Foundation, Award ID: 854391 and JLW by P01 HL147835.

## Author contributions

MDG performed *in vitro* and *in vivo* experiments, conceived the experiments, and helped to write the manuscript; VP performed CLP experiments and the analyses related to *in vivo* experiments; PJT prepared oxPAPC; RS performed the bioinformatics analyses; AC identified and isolated samples from septic patients; AGC performed CLP experiments; PLSMG performed the MRSA experiments; LP and FM analyzed oxPL levels in the BALs; JLW provided transgenic mice and gave advice; JC helped to perform and analyze the experiments with human samples; JRS provided oxPAPC and gave advices; IZ conceived and supervised the experiments, and wrote the paper.

## Supplementary Figure Legends

**Supplementary Figure 1.**
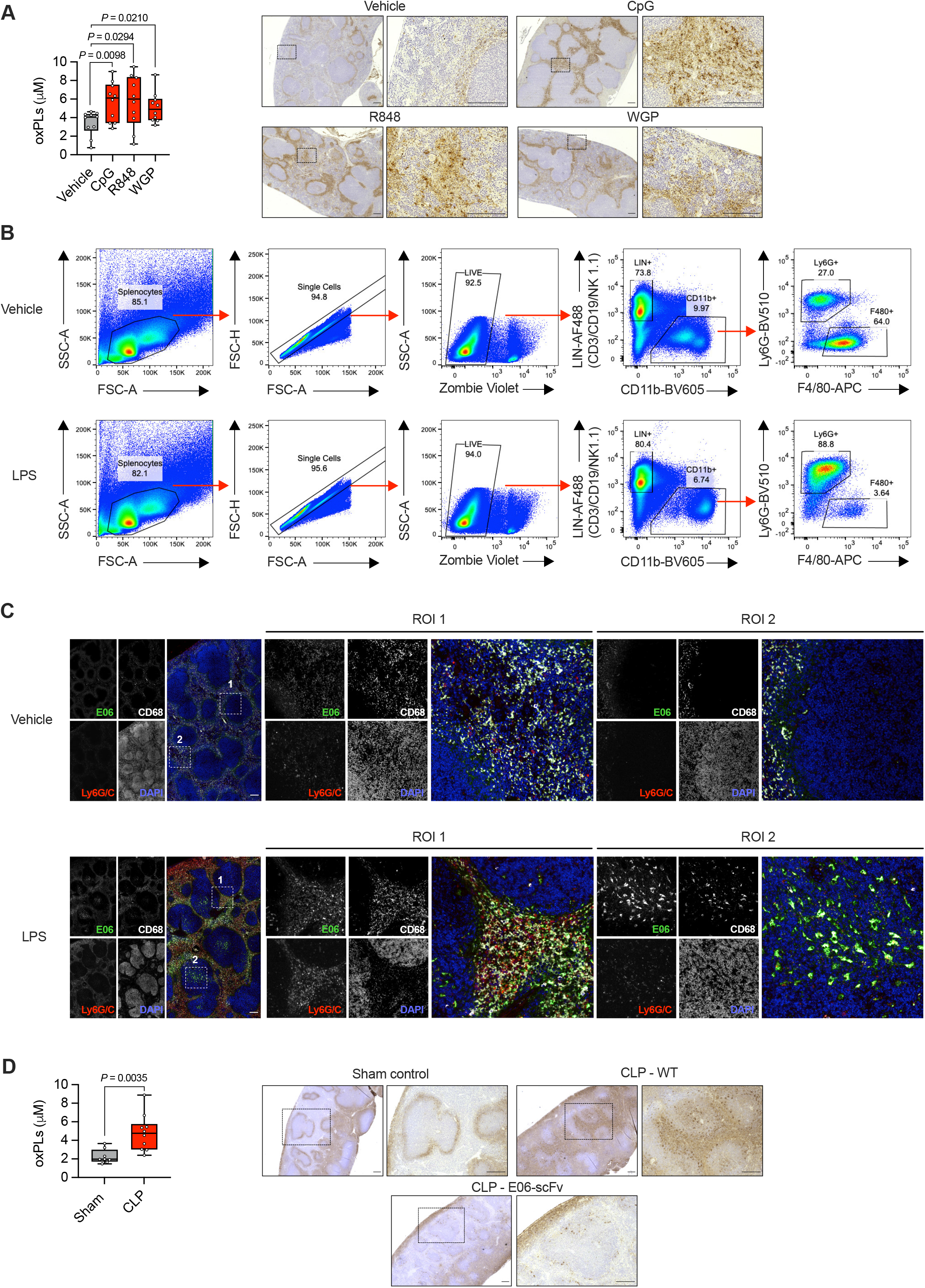
**A**) Mice were i.p. injected with vehicle (saline), CpG (15 mg Kg^−1^), R848 (10 mg Kg^−1^) or WGP (15 mg Kg^−1^). oxPLs were quantified in serum after 8h (*left*) or in spleen after 24h (*right*). *n* = 10 mice per group. Statistical significance was calculated using two-tailed *t* test. Spleens were immunostained with E06 antibody. Representative photomicrographs are shown. Scale bar = 200 μm. **B, C**) Mice were treated with vehicle (saline) or LPS (2 mg Kg^−1^) for 24h and splenic cell populations were analyzed for oxPLs accumulation using flow cytometry (**B**, gate strategy) or (**C**) fluorescent immunohistochemistry. Representative photomicrographs are shown. Scale bar = 200 μm. **D**) Mice were subjected to CLP. oxPLs were quantified in serum after 8h (*left*) or in spleen after 24h (*right*). *n* = 8 (sham control) *n* = 10 (CLP) mice per group. Statistical significance was calculated using two-tailed *t* test. Spleens were immunostained with E06 antibody. Representative photomicrographs are shown. Scale bar = 200 μm.

**Supplementary Figure 2.**
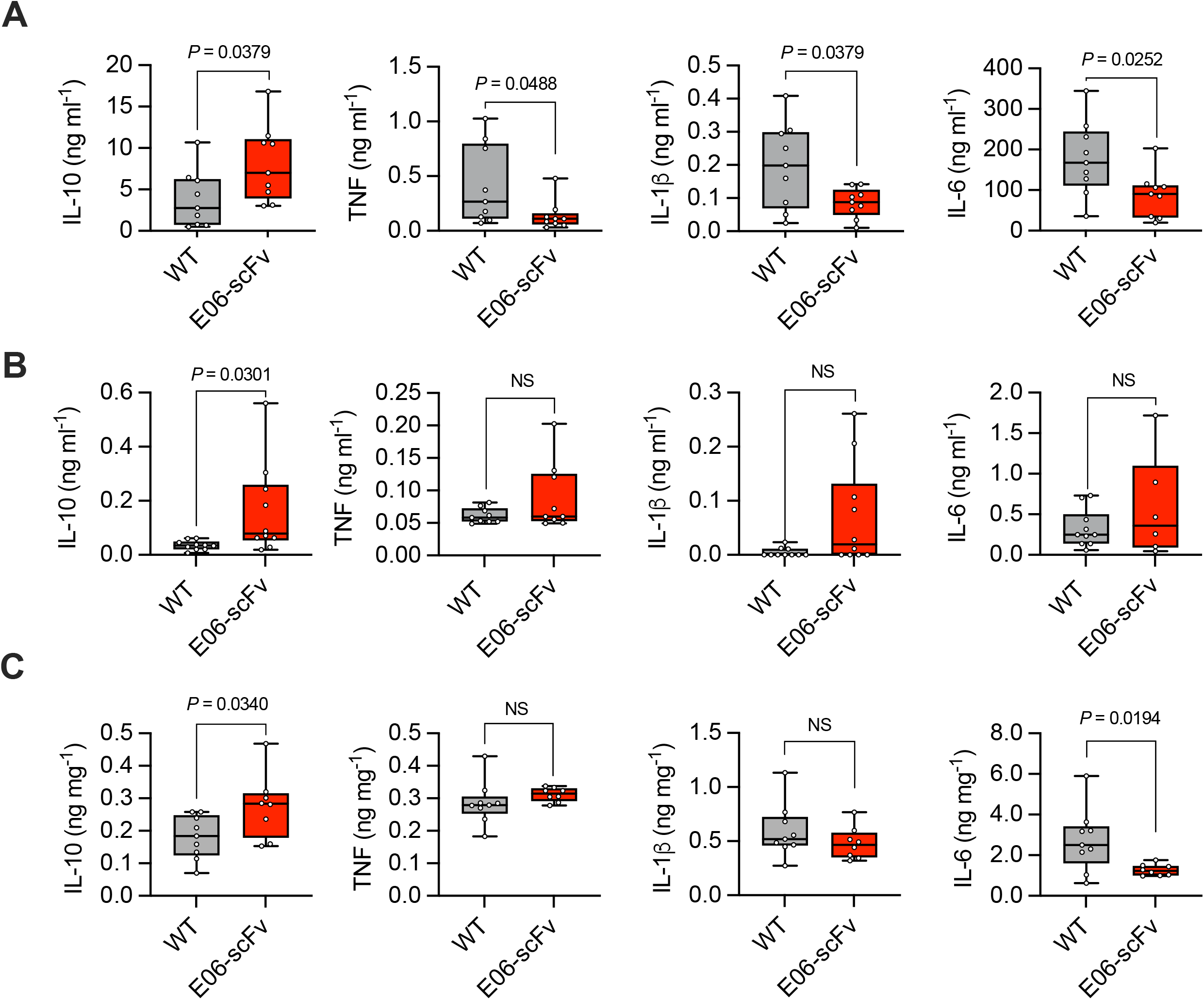
**A**) WT and E06-scFv mice (*n* = 9) were subjected to CLP. The indicated cytokines were quantified after 8h from serum. Graph shows means ± SD. Statistical significance was calculated using two-tailed *t* test. **B**) WT and E06-scFv mice (n = 10) were injected with MRSA (1 × 10^8^ CFU/mouse) and the indicated cytokines were measured from serum 24h post-infection. Graph shows means ± SD. Statistical significance was calculated using Mann-Whitney Test. **C**) WT (*n* = 9) and E06-scFv (*n* = 8) mice were intratracheally daily administered with poly(I:C) (2.5 mg Kg^−1^). At day 6, the indicated cytokines were measured from lung homogenates. Graph shows means ± SD. Statistical significance was calculated using two-tailed *t* test.

**Supplementary Figure 3.**
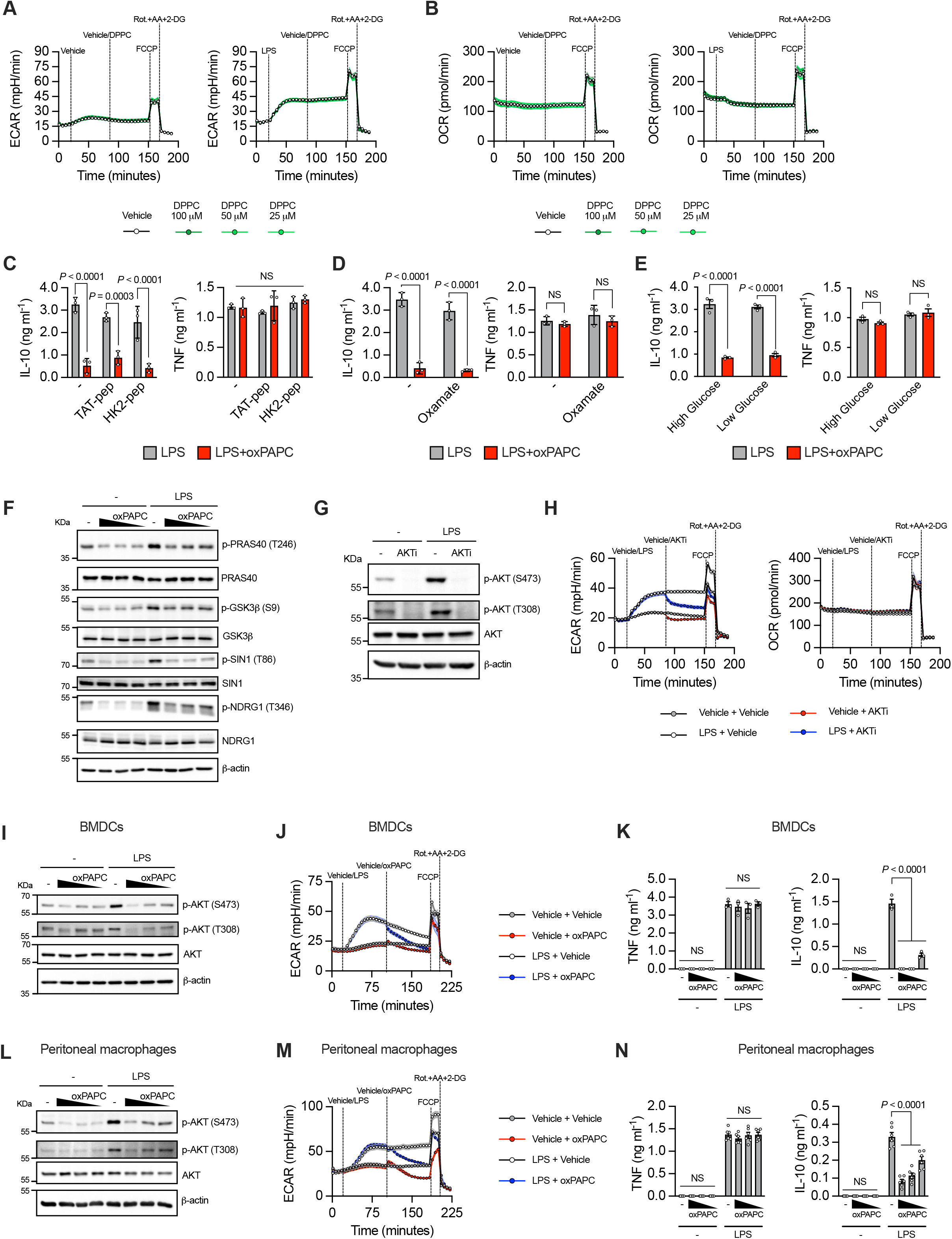
**A, B**) Real-time changes in ECAR (**A**) and OCR (**B**) of BMDMs untreated or treated with LPS (1 μg ml^−1^) and then challenged with different doses of DPPC, as indicated. *n* = 6, graphs are representative of three independent experiments and show means ± SEM. **C-E**) BMDMs were incubated with TAT-peptide (10 μM), TAT-HK2-peptide (10 μM) (**C**), oxamate (30 mM) (**D**) or seed in high glucose (25 mM) and low glucose (1 mM) medium (**E**) for 30 minutes. The cells were primed with LPS and then stimulated with oxPAPC (100 μg ml^−1^). IL-10 and TNF production was quantified by ELISA after 18h. *n* = 3, graphs are representative of three independent experiments and show means ± SEM. Statistical multiple comparisons were calculated by two-way ANOVA and Sidak’s multiple comparisons test. **F**) BMDMs were primed, or not, with LPS and then stimulated with oxPAPC (100, 50 or 25 μg ml^−1^). The indicated AKT targets and downstream kinases were analyzed after 1h by immunoblotting. Data are representative of three independent experiments. **G**) BMDMs were primed, or not, with LPS and then treated with AKTi (10 μM). AKT phosphorylation was analyzed after 1h by immunoblotting. Data are representative of three independent experiments. **H**) Real-time changes in ECAR (*left*) and OCR (*right*) of BMDMs untreated or treated with LPS (1 μg ml^−1^) and then challenged with AKTi (10 μM), as indicated. *n* = 6, graphs are representative of three independent experiments and show means ± SEM. **I**) Murine GM-CSF-derived DCs (BMDCs) were primed, or not, with LPS and treated or not with oxPAPC (100, 50 or 25 μg ml^−1^) for 1h. AKT phosphorylation was assessed by immunoblot. Data are representative of three independent experiments **J**) ECAR of BMDCs, treated as indicated, was measured using a Seahorse analyzer. oxPAPC was used at 100 μg ml^−1^. Data are representative of three independent experiments. **K**) BMDCs were treated as in **I** and TNF and IL-10 release was quantified by ELISA after 18h. *n* = 3, graphs are representative of three independent experiments and show means ± SEM. Statistical multiple comparisons were calculated by two-way ANOVA and Dunnett’s test. **L**) Murine peritoneal macrophages were primed, or not, with LPS and treated or not with oxPAPC (50, 12 or 12.5 μg ml^−1^) for 1h. AKT phosphorylation was assessed by immunoblot. Data are representative of three independent experiments. **M**) ECAR murine peritoneal macrophages, treated as indicated, was measured using a Seahorse analyzer. oxPAPC was used at 50 μg ml^−1^. Data are representative of three independent experiments. **N**) Murine peritoneal macrophages were treated as in **L** and TNF and IL-10 release was quantified by ELISA after 18h. *n* = 6, graphs are representative of three independent experiments and show means ± SEM. Statistical multiple comparisons were calculated by two-way ANOVA and Dunnett’s test.

**Supplementary Figure 4.**
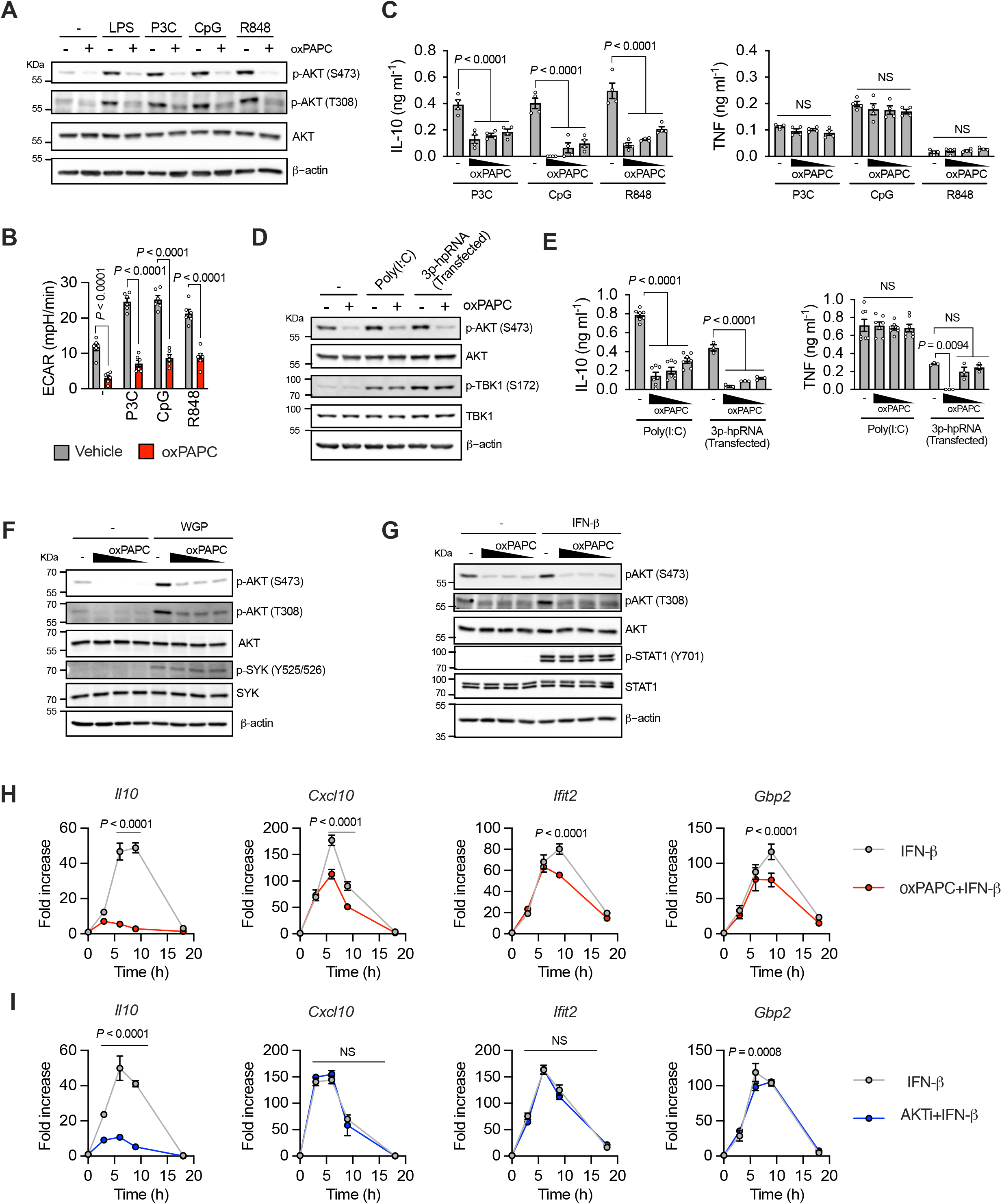
**A**) BMDMs were primed, or not, with LPS (1 μg ml^−1^), P3C (1 μg ml^−1^), CpG (5 μM) or R848 (1 μg ml^−1^) and then stimulated with oxPAPC (100, 50 or 25 μg ml^−1^). AKT phosphorylation was analyzed after 1h by immunoblotting. Data are representative of three independent experiments. **B**) ECAR of BMDMs, treated as indicated, was measured using a Seahorse analyzer. oxPAPC was used at 100 μg ml^−1^. *n* = 6, data are representative of three independent experiments and show means ± SEM. Statistical significance was calculated using two-way ANOVA and Sidak’s multiple comparisons test. **C**) BMDMs were treated as in (**A**) and IL-10 and TNF release was quantified by ELISA after 18h. *n* = 4, graphs are representative of three independent experiments and show means ± SEM. Statistical multiple comparisons were calculated by two-way ANOVA and Dunnett’s test. **D**) BMDMs were primed, or not, with poly(I:C) (10 μg ml^−1^) or transfected with 3p-hpRNA/LyoVec complexes (1 μg ml^−1^) and then stimulated with oxPAPC (50 μg ml^−1^). AKT phosphorylation was analyzed after 1h by immunoblotting. Data are representative of three independent experiments. **E**) BMDMs were primed with poly(I:C) (*n* = 7) or 3p-hpRNA/LyoVec (*n* = 3) as in (**D**) and then stimulated with oxPAPC (100, 50 or 25 μg ml^−1^). IL-10 and TNF release was quantified by ELISA after 18h. Graphs are representative of three independent experiments and show means ± SEM. Statistical multiple comparisons were calculated by two-way ANOVA and Dunnett’s test. **F**) BMDMs were pre-treated with GM-CSF. After 16h, cells were primed, or not, with WGP (1 μg ml^−1^) and then stimulated with oxPAPC (100, 50 or 25 μg ml^−1^). AKT phosphorylation was analyzed after 1h by immunoblotting. Data are representative of three independent experiments. **G**) BMDMs were treated with oxPAPC (100, 50 or 25 μg ml^−1^) for 30 minutes and then stimulated, or not, with IFNβ (10 ng ml^−1^) for 1h. AKT and STAT1 phosphorylation were analyzed after 1h by immunoblotting. Data are representative of three independent experiments. **H-I**) BMDMs were treated with vehicle or oxPAPC (100 μg ml^−1^) (**H**) or AKTi (10 μM) (**I**) for 30 minutes and then stimulated with IFNβ (10 ng ml^−1^). The indicated transcripts were analyzed by qPCR at the indicated time points. *n* = 3, data are representative of three independent experiments and show means ± SEM. Statistical significance was calculated using two-way ANOVA and Sidak’s multiple comparisons test.

**Supplementary Figure 5.**
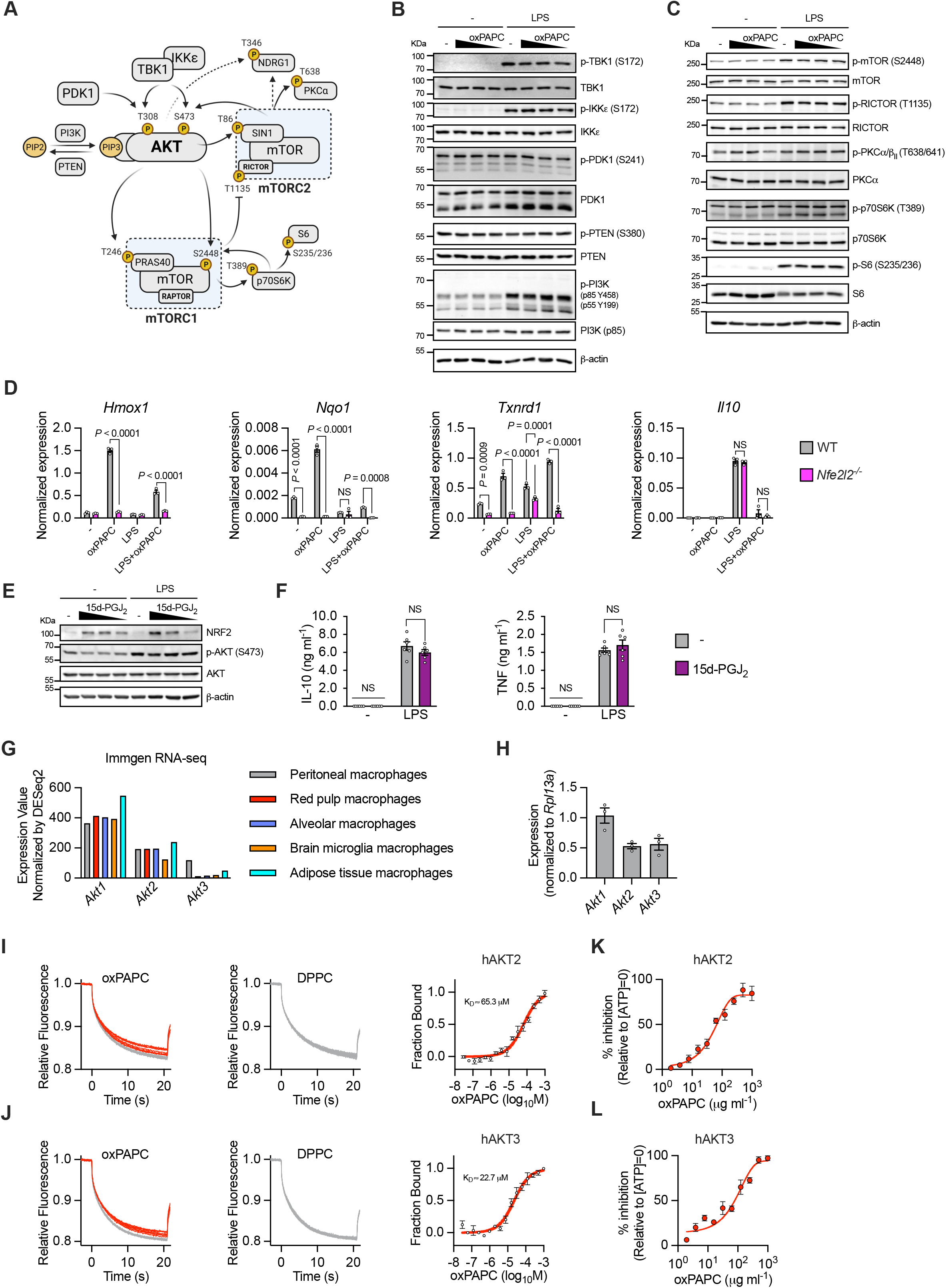
**A**) Schematic shows upstream and downstream kinases in the AKT signaling network. **B**) BMDMs were primed, or not, with LPS and then stimulated with oxPAPC (100, 50 or 25 μg ml^−1^). Phosphorylation of upstream AKT regulators (TBK1, IKKε, PDK1, PTEN and PI3K) was analyzed by immunoblot. Images are representative of three independent experiments. **C**) BMDMs were primed, or not, with LPS (1 μg ml^−1^) and then stimulated with oxPAPC (100, 50 or 25 μg ml^−1^). The phosphorylation of the indicated mTORC1/2-associated proteins was analyzed after 1h by immunoblotting. Data are representative of three independent experiments. **D**) WT and *Nfe2l2^−/−^* BMDMs were primed, or not, with LPS (1 μg ml^−1^) and treated or not with oxPAPC (100 μg ml^−1^) for 1h. The indicated transcripts were analyzed by qPCR. *n* = 3, data are representative of three independent experiments and show means ± SEM. Statistical significance was calculated using two-way ANOVA and Sidak’s multiple comparisons test. **E-F**) BMDMs were primed, or not, with LPS (1 μg ml^−1^) and treated or not with 15d-PGJ_2_ (20, 10 or 1 μM in **E** and 10 μM in **F**) for 1h (**E**) or 24h (**F**). AKT phosphorylation and NRF2 accumulation were assessed by immunoblot (**E**). Data are representative of three independent experiments. IL-10 and TNF release was quantified by ELISA (**F**). *n* = 6, graphs are representative of three independent experiments and show means ± SEM. Statistical multiple comparisons were calculated by two-way ANOVA and Sidak’s test. **G**) Expression of *Akt* isoforms (RNA-seq, Immgen.org database) in the indicated murine macrophage populations. **H**) Normalized expression of *Akt1*, *Akt2* and *Akt3* in BMDMs, analyzed by qPCR. *n* = 3, data are representative of three independent experiments and show means ± SEM. **I, J**) Microscale thermophoresis (MST) analysis of oxPAPC interactions with AKT2 (**I**) and AKT3 (**J**). Traces of fluorescently labeled human recombinant AKT incubated with oxPAPC (1000, 500, 250,125, 62.5, 31.25, 15.62, 7.8, 3.9, 1.9, 0.97, 0.488, 0.24, 0.12, 0.0061 and 0.030 μM) or DPPC (500, 250,125, 62.5, 31.25, 15.62, 7.8, 3.9, 1.9, 0.97, 0.488, 0.24, 0.12, 0.0061 and 0.030 μM). oxPAPC-AKT binding curve was derived from the quantification of normalized fluorescence changes. *n* = 3, graph shows means ± SD. Images are representative of three independent experiments. **K, L**) Active human recombinant AKT2 (**K**) or AKT3 (**L**) were incubated with oxPAPC and a kinase-specific FRET-peptide substrate (Z-’LYTE). Kinase inhibition was measured as the ability of lipid to block substrate phosphorylation. *n* = 4, graph shows means ± SD. Data are representative of three independent experiments.

**Supplementary Figure 6.**
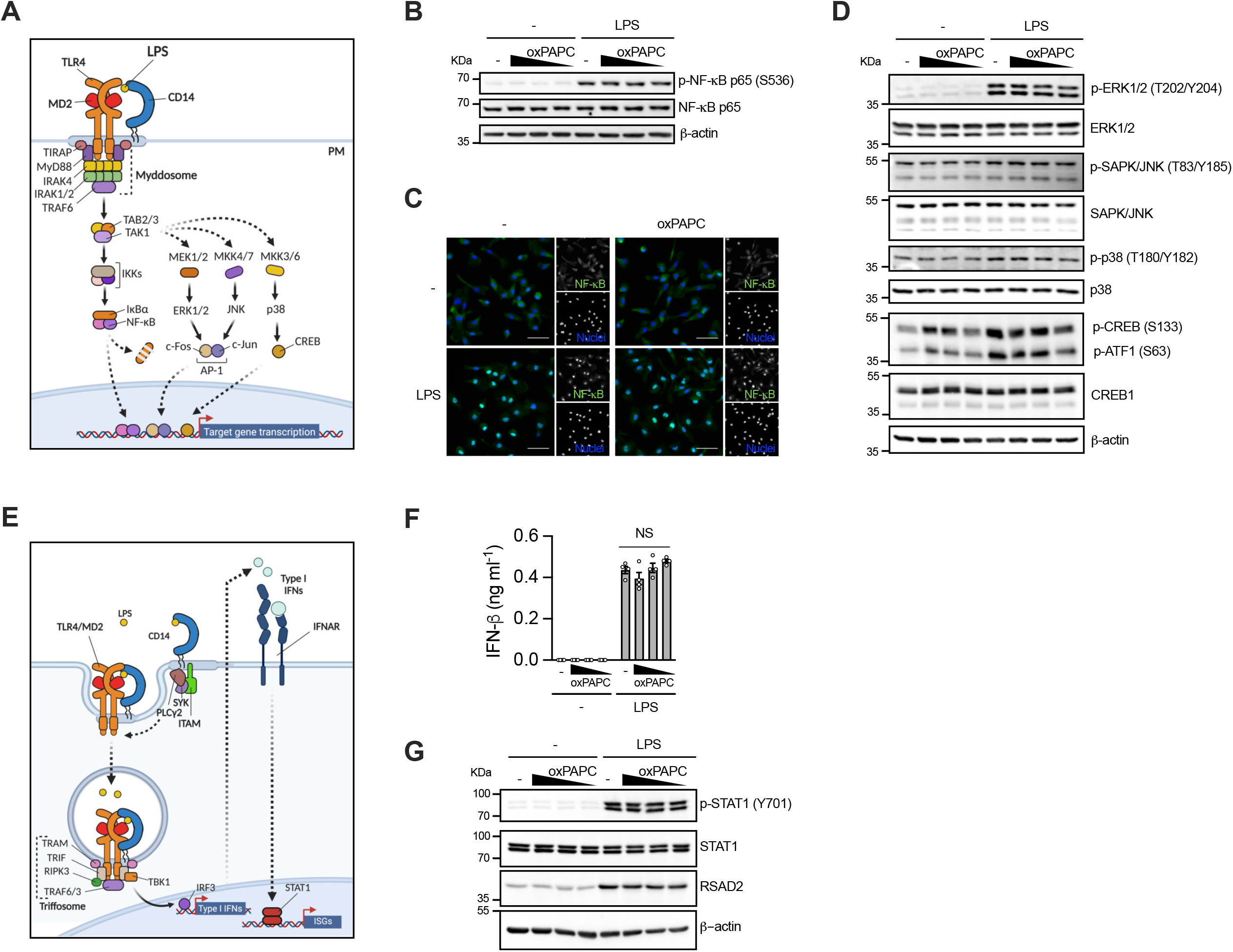
**A**) Schematic shows MyD88-dependent TLR4 downstream signaling. **B-D**) BMDMs were primed, or not, with LPS (1 μg ml^−1^) and treated or not with oxPAPC (100 μg ml^−1^ in **B**, **D** and 100, 50 or 25 μg ml^−1^ in **B, D** and 100 μg ml^−1^ in **C**) for 1h. NF-κB (**B**) and MAPK pathway activation (**D**) were assessed by immunoblot. Images are representative of three independent experiments. NF-κB nuclear translocation was analyzed by immunofluorescence (**C**). Scale bar 50 μm. Data are representative of three independent experiments. **E**) Schematic shows TRIF-dependent TLR4 downstream signaling. **F, G**) BMDMs were primed, or not, with LPS (1 μg ml^−1^) and treated or not with oxPAPC (100, 50 or 25 μg ml^−1^). IFN-β production (**F**) was quantified by ELISA after 24h. *n* = 4, data are representative of three independent experiments and show means ± SEM. Statistical multiple comparisons were calculated by two-way ANOVA and Tukey’s test. Type I IFN pathway activation was assessed by immunoblot (**G**). Data are representative of three independent experiments.

**Supplementary Figure 7.**
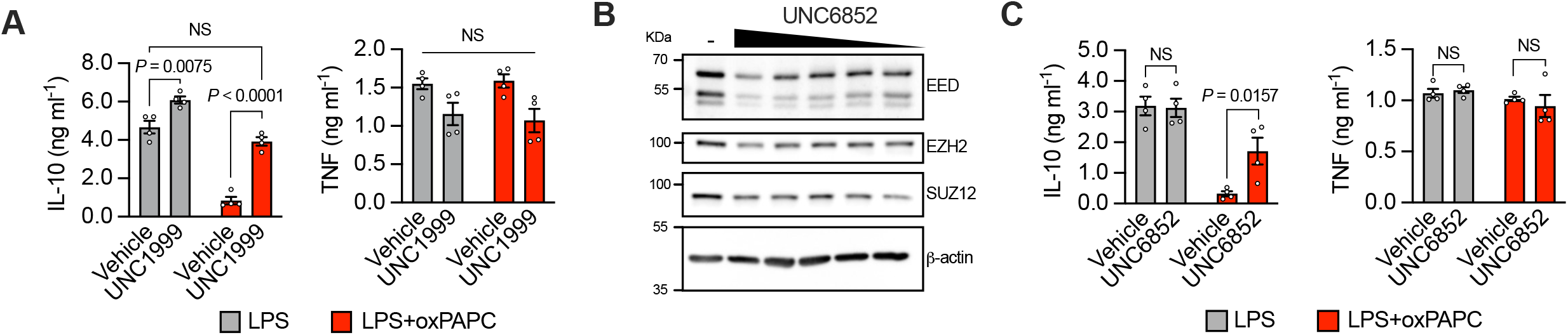
**A**) BMDMs were incubated with UNC1999 (5 μM) for 30 minutes, primed with LPS and then stimulated with oxPAPC (100 μg ml^−1^). IL-10 and TNF production was quantified by ELISA after 18h. *n* = 4, graphs are representative of three independent experiments and show means ± SEM. Statistical multiple comparisons were calculated by two-way ANOVA and Tukey’s test. **B**) BMDMs were treated with UNC6852 (50, 25, 12.5, 6.25, 3 and 1.5 μM) for 24h and the indicated components of PRC2 complex were quantified by immunoblot. Image is representative of three independent experiments. **C**) BMDMs were incubated with UNC6852 (50 μM) for 24h, primed with LPS and then stimulated with oxPAPC (100 μg ml^−1^). IL-10 and TNF production was quantified by ELISA after 18h. *n* = 4, graphs are representative of three independent experiments and show means ± SEM. Statistical multiple comparisons were calculated by two-way ANOVA and Tukey’s test.

## Methods

### Mouse strains and murine cell cultures

C57BL/6J (Jax 000664), *Tlr4^−/−^* (Jax 029015), *Tlr2^−/−^* (Jax 004650), *Cd14^−/−^* (Jax 003726), *Cd36^−/−^* (Jax 019006), *Nfe2l2^−/−^* (Jax 017009) mice were acquired from The Jackson Laboratory. E06-scFv animals on C57BL/6J background were provided by J.L. Witztum (University of California San Diego). Macrophages were differentiated from bone marrow using Dulbecco’s modified Eagle medium (DMEM, Thermo Fisher Scientific, Cat# 11965092), with 30% L929-M-CSF (BMDMs) or 10% B16-GM-CSF supernatant (GM-derived phagocytes), 10% fetal bovine serum (FBS), 2mM L-glutamine (Thermo Fisher Scientific, Cat# 25030081) and 100U/100μg ml^−1^ penicillin/streptomycin (Thermo Fisher Scientific, Cat# 15140122). Primary peritoneal macrophages were obtained by peritoneal lavage of mice treated intraperitoneally with 4% thioglycolate (Sigma-Aldrich, Cat# T9032) for 4 days. Prior to stimulations, cultured cells were rinsed with phosphate-buffered saline (PBS) and re-plated in DMEM supplemented with 10% FBS, 2mM L-glutamine and 100U/100μg ml^−1^ penicillin/streptomycin at a concentration of 1 × 10^6^ cells ml^−1^. In all experiments requiring LPS priming, LPS was used at the concentration of 1 μg ml^−1^ for 3 hours, unless otherwise stated.

### Human phagocytes cell cultures

Human phagocytes were isolated or differentiated from collars of blood received from Boston Children’s Hospital healthy donors. Briefly, blood was diluted 1:2 in PBS and peripheral blood mononuclear cells (PBMCs) were isolated using Histopaque (Sigma, Cat# 1077-1) gradient. Monocytes were positively selected from PBMCs with CD14 MicroBeads (Miltenyi Biotec, Cat# 130-050-201). Mo-DCs and macrophages were differentiated from monocytes in Roswell Park Memorial Institute (RPMI) 1640 medium (Thermo Fisher Scientific, 61870127) supplemented with 10% FBS in presence of GM-CSF (20 ng ml^−1^, BioLegend Cat# 572903) and IL-4 (20 ng ml^−1^, BioLegend Cat# 574004) (Mo-DCs) or M-CSF (50 ng ml^−1^, BioLegend Cat# 574804) and IL-4 (20 ng ml^−1^) (macrophages) for 7 days. Prior to stimulations, cultured cells were rinsed with PBS and re-plated in RPMI 1640 supplemented with 10% FBS, 2mM L-glutamine and 100U/100μg ml^−1^ penicillin/streptomycin at a concentration of 1 × 10^6^ cells ml^−1^. In all experiments with human cells requiring LPS priming, LPS was used at the concentration of 0.1 μg ml^−1^ for 3 hours, unless otherwise stated.

### Sepsis patient cohort and clinical characteristics

The Third International Consensus Definitions for Sepsis and Septic Shock were used to identify patients with sepsis [31]. Patients with sepsis had an identified infection with > 2 point increase in the Pediatric Sequential Organ Failure Assessment score at the time of study enrollment compared to their pre-admission score. All patients were admitted to the critical care unit, with blood obtained within 48 hours of admission. This study was approved by the institutional review board of Boston Children’s Hospital.

### Clinical samples for oxPLs measurement in BAL

BALs were obtained from COVID-19 and control patients previously described in[56]. Briefly, BALs from SARS-CoV-2 positive patients (27) hospitalized in the Intensive Care Unit (ICU) were collected from September to November 2020 at Luigi Sacco Hospital (Milan, Italy). Control samples, sarcoidosis patients (10) and lung transplant patients (10), were collected by IRCCS Policlinico San Matteo Foundation (Pavia, Italy). None of the control patients were diagnosed a respiratory infection.

### Stimuli and reagents

oxPAPC and oxPAPE-N-Biotin were custom made starting from PAPC (Avanti Polar Lipids, Cat# 850459) and PAPE (Avanti Polar Lipids, Cat# 850759). DPPC (Cat# 850355) was purchased from Avanti Polar Lipids. 15d-PGJ_2_ (Cat# 18570) and 2-deoxy-D-Glucose (2-DG, Cat# 14325) were purchased from Cayman Chemicals. *Escherichia coli* LPS (Serotype O555:B5, TLRgrade, Cat# ALX-581-013-L002) was purchased from Enzo Life Sciences. Pam3CSK4 (Cat# tlrl-pms), CpG ODN 1826 (Cat# tlrl-1826), R848 (Cat# tlrl-r848), poly(I:C) (Cat# tlrl-pic-5), 5’ triphosphate hairpin RNA (3p-hpRNA, Cat# tlrl-hprna), LyoVec (Cat# lyec-1) and dispersible whole glucan particles (WGP, Cat# tlrl-wgp) were purchased from InvivoGen. Recombinant murine IFN-β1 (Cat# 581304) was purchased from BioLegend. AKT inhibitor VIII (Cat# A6730), sodium oxamate (Cat# O2751), FCCP (Cat# C2920), oligomycin A (OM, Cat# 75351), rotenone (Cat# R8875) and Antimycin A (Cat# A8674) were purchased from Sigma-Aldrich. GSK343 (Cat# T6059), UNC6852 (Cat# T13954) and UN1999 (Cat# T3057) were purchased from TargetMol Chemicals. Peptides TAT-HKII (MIASHMIACLFTELN(β-Ala)GYGRKKRRQRRG-amide) and TAT (GYGRKKRRQRRG-amide) were custom-made by GenScript.

### oxPAPE-N-Biotin synthesis

A solution of PAPE in dry dichloromethane was added dropwise to a magnetically stirred solution of biotin, N,N’-dicyclohexylcarbodiimide (DCC) and 4-dimethylaminopyridine (DMAP) in dry dichloromethane under argon at room temperature. The solution was mixed with a magnetic stirrer under argon for 12-24h at room temperature. oxPAPE-N-Biotin was directly purified with a syringe filter (PES 0.45 um) and SPE kit.

### Lipid oxidation

PAPC and PAPE-N-Biotin were oxidized using a method similar to that previously reported [57]. Briefly, phospholipids were transferred to clean borosilicate tubes in 0.5 mg aliquots in chloroform, dried, and oxidized for 24–72h, while monitoring oxidation by flow injection on an electrospray ionization (ESI) instrument (LCQ Fleet, Thermo Fisher Scientific). Lipid concentrations were analyzed by phosphorus assay.

### Seahorse metabolic analysis

The ECAR and OCR were measured with a Seahorse XFe96 Extracellular Flux Analyzer. Phagocytes (8 × 10^4^ per well) were seeded in a Seahorse 96-well plate in DMEM complete medium (murine cells) or RPMI complete medium (human cells). After 4h, cells were washed and incubated in the Seahorse Assay Medium (Agilent Technologies, Seahorse XF DMEM medium, Cat# 103575-100, or Seahorse XF RPMI medium, Cat# 103576-100) supplemented with 10 mM glucose and 2 mM glutamine at 37 °C for 45 min. The OCR and ECAR were measured under basal conditions and after injection of LPS (1 or 0.1 μg ml^−1^), oxPAPC, DPPC, FCCP (1.5 μM) and rotenone (0.5 μM) plus antimycin A (0.5 μM) plus 2-DG (50 mM) (rot. + AA + 2-DG), as indicated in the figures.

### Gene expression analysis

RNA was isolated from cell cultures using Direct-zol RNA Miniprep Kit (Zymo Research, Cat# R2053). Purified RNA was analyzed for gene expression on a CFX384 real-time cycler (Bio-Rad Laboratories) using Power SYBR Green RNA-to-CT 1-Step Kit (Thermo Fisher Scientific, Cat# 4389986) with predesigned primers (Sigma-Aldrich, KiCqStart SYBR Green Primers) specific for murine *Il10*, *Cxcl10*, *Ifit2*, *Gbp2*, *Hmox1*, *Nqo1*, *Txnrd1* and *Rpl13a* (housekeeping). In order to analyze *Akt* isoforms expression, TaqMan RNA-to-CT 1-Step Kit (Thermo Fisher Scientific, Cat# 4392938) was used in association with the following probes: *Akt1* (Thermo Fisher Scientific, Cat# Mm01331626_m1), *Akt2* (Thermo Fisher Scientific, Cat# Mm05804787_gH), *Akt3* (Thermo Fisher Scientific, Cat# Mm00442194_m1) and *Rpl13a* (Thermo Fisher Scientific, Cat# Mm05910660_g1).

### Measurement of cytokines by ELISA

Cytokine production was quantified from cell supernatants or serum using the following ELISA kits: ELISA MAX Deluxe Set Mouse IL-10 (BioLegend, Cat# 431414), ELISA MAX Deluxe Set Human IL-10 (BioLegend, Cat# 430604), ELISA MAX Deluxe Set Mouse TNF-α (BioLegend, Cat# 430915), ELISA MAX Deluxe Set Human TNF-α (BioLegend, Cat# 430204), ELISA MAX Deluxe Set Mouse IL-6 (BioLegend, Cat# 431315) and ELISA MAX Deluxe Set Mouse IL-1β (BioLegend, Cat# 432604).

### Immunoblotting

For western blotting, phagocytes (1 × 10^6^) were treated as indicated and subsequently lysed in 100 μl RIPA buffer (Boston BioProducts, Cat# BP-116TX) added with protease and phosphatase inhibitors (Thermo Fisher Scientific, Halt Protease and Phosphatase Inhibitor Cocktail, Cat# 78440). Total protein concentrations were determined using BCA protein assay kit (Genesee Scientific, Cat# 18-440). Proteins were separated using SDS-PAGE electrophoresis and then transferred to polyvinylidene difluoride (PVDF) membranes. After 1h incubation with blocking buffer (5% milk or BSA in Tris-buffered saline (TBS) with 0.05% Tween-20), membranes were probed with the following primary antibodies: phospho-AKT (Ser473) (1:1000, Cell Signaling Technology, Cat# 4060), phospho-AKT (Thr308) (1:1000, Cell Signaling Technology, Cat# 13038), AKT (pan) (1:1000, Cell Signaling Technology, Cat# 4691), β-actin (1:3000, Sigma–Aldrich, Cat# A5441), phospho-SIN1 (Thr86) (1:1000, Cell Signaling Technology, Cat# 14716), SIN1 (Proteintech Group, Cat# 15463-1-AP), phospho-NDRG1 (Thr346) (1:1000, Cell Signaling Technology, Cat# 5482), NDRG1 (1:1000, Abcam, Cat# ab37897), phospho-PRAS40 (Thr246) (1:1000, Cell Signaling Technology, Cat# #13175), PRAS40 (Cell Signaling Technology, Cat# #2610), phospho-GSK3β (Ser9) (1:1000, Cell Signaling Technology, Cat# #5558), GSK3β (Proteintech Group, Cat# 22104-1-AP), phospho-TBK1 (Ser172) (1:1000, Cell Signaling Technology, Cat# 5483), TBK1 (1:1000, Cell Signaling Technology, Cat# 38066), phospho-IKKε (Ser172) (1:1000, Sigma-Aldrich, Cat# 06-1340), IKKε (1:1000, Cell Signaling Technology, Cat# 3416), phospho-PDK1 (Ser241) (1:1000, Cell Signaling Technology, Cat# #3438), PDK1 (1:1000, Cell Signaling Technology, Cat# 3062), phospho-PTEN (Ser380) (1:1000, Cell Signaling Technology, Cat# 9551), PTEN (1:1000, Cell Signaling Technology, Cat# 9559), phospho-PI3K p85 (Tyr458)/p55 (Tyr199) (1:1000, Cell Signaling Technology, Cat# 4228), PI3K p85 (1:1000, Cell Signaling Technology, Cat# 4257), phospho-SYK (Tyr525/526) (1:1000, Cell Signaling Technology, Cat# 2710), SYK (1:1000, Cell Signaling Technology, Cat# 13198), phospho-mTOR (Ser2448) (1:1000, Cell Signaling Technology, Cat# 5536), mTOR (1:1000, Cell Signaling Technology, Cat# 2983), phospho-RICTOR (Thr1135) (1:1000, Cell Signaling Technology, Cat# 3806), RICTOR (1:1000, Cell Signaling Technology, Cat# 2114), phospho-PKCα/β_II_ (Thr638/641) (1:1000, Cell Signaling Technology, Cat# 9375), PKCα (Proteintech Group, Cat# 21991-1-AP), phospho-p70S6K (Thr389) (1:1000, Cell Signaling Technology, Cat# 9234), p70S6K (1:1000, Cell Signaling Technology, Cat# 34475), phospho-S6 (Ser235/Ser236) (1:500, BioLegend, Cat# 608602), S6 (1:500, BioLegend, Cat# 691802), phospho-STAT1 (Tyr701) (1:1000, Becton, Dickinson and Company, Cat# 612133), STAT1 (1:1000, Cell Signaling Technology, Cat# 14994), RSAD2 (1:1000, BioLegend, custom made 91736), NRF2 (1:1000, Cell Signaling Technology, Cat# 12721), phospho-ERK1/2 (Thr202/Tyr204) (1:1000, Cell Signaling Technology, Cat# 9106), ERK1/2 (1:1000, Cell Signaling Technology, Cat# 9102), phospho-p38 (Thr180/Tyr182) (1:1000, Cell Signaling Technology, Cat# 4511), p38 (1:500, BioLegend, Cat# 622403), phospho-SAPK/JNK (Thr183/Tyr185) (1:1000, Cell Signaling Technology, Cat# 4668), SAPK/JNK (1:1000, Cell Signaling Technology, Cat# 9252), phospho-CREB (Ser133) (1:1000, Cell Signaling Technology, Cat# 9198), CREB1 (1:2000, Proteintech Group, Cat# 12208-1-AP), phospho-NF-κB (Ser536) (1:1000, Cell Signaling Technology, Cat# 3033), NFκB p65 (1:1000, Cell Signaling Technology, Cat# 6956), EED (1:1000, Cell Signaling Technology, Cat# 85322), EZH2 (1:1000, Cell Signaling Technology, Cat# 5246) and SUZ12 (1:1000, Cell Signaling Technology, Cat# 3737).

After incubation with the specific HRP-conjugated antibodies anti-rabbit (1:10000, Thermo Fisher Scientific, Cat# G-21234), anti-mouse (1:10000, Jackson ImmunoResearch, Cat# 715-035-151) or anti-goat (1:10000, Santa Cruz Biotechnology, Cat# sc-2354), the immunoreactive bands were detected by iBright FL100 Imaging System (Thermo Fisher Scientific).

### Measurement of total oxPLs by ELISA

The concentration of total immunodetectable oxPLs in mouse and human samples (serum, BAL or lung homogenate) was measured using a modification of a competitive ELISA technique, as previously described [13]. Briefly, white immunoplates (Thermo Fisher Scientific, Cat# 436110) were coated with an IgG form of the E06 antibody (Absolute Antibody, Cat# Ab02746-23.0) and left overnight at 4°C. Plates were blocked with 1% fatty acid free BSA for 1 hour and then diluted serum samples were added in triplicate to the wells. Next, Phosphorylcholine Keyhole Limpet Hemocyanin (PC-KLH, LGC Biosearch Technologies, Cat# PC-1013-5) was applied at a concentration of 100 ng per well. After 1h incubation, alkaline phosphatase conjugated anti-KLH antibody (Rockland Inc, Cat# 600–405-466) was added to detect the bound PC-KLH, using Lumi-Phos 530 (Lumigen, Cat# P-501). The luminescence emitted was measured using a SpectraMax i3x plate reader (Molecular Devices). In order to calculate the concentration of oxPLs in the samples, each plate included a standard curve of oxPAPC.

### Flow Cytometry

Spleens collected from the indicated mice were mechanically dissociated and filtered to obtain a cell suspension. After red blood cell lysis, dead cells were stained with Zombie Violet (BioLegend, Cat# 423114) and cellular surface markers were stained with the following fluorophore-conjugated anti-mouse antibodies: Alexa Fluor 488 CD3 (BioLegend, Cat# 100210), Alexa Fluor 488 CD19 (BioLegend, Cat# 115521), Alexa Fluor 488 NK1.1 (BioLegend, Cat# 108718), BV605 CD11b (BioLegend, Cat# 101257), BV510 Ly6G (BioLegend, Cat# 127633), APC F4/80 (BioLegend, Cat# 123116). Cells were fixed (Fixation buffer, BioLeged, Cat# 420801), permeabilized (Intracellular Staining Permeabilization Wash Buffer, BioLegend, Cat# 421002) and incubated for 30 minutes with TruStain FcX™ PLUS (anti-mouse CD16/32) antibody (BioLegend, Cat# 156604). Immunodetectable oxPLs were analyzed using custom-made PE-conjugated E06 rabbit antibody (Absolute antibody) and PE-conjugated isotype control (rabbit IgG, Cell Signaling Technology, Cat# 5742S).

To analyze the phosphorylation status of EZH2, BMDMs were treated as indicated, fixed and permeabilized using True-Phos Perm Buffer (BioLegend, Cat# 425401). Cells were incubated with phospho-EZH2 (S21) (Affinity Biosciences, Cat# AF3822) or rabbit IgG isotype control (BioLegend, Cat# 910801) followed by PE-conjugated anti-rabbit antibody (BioLegend, Cat# 406421) staining.

Samples were acquired with LSRFortessa Cell Analyzer (Becton, Dickinson and Company) and analyzed by FlowJo v10.9 (Becton, Dickinson and Company).

### Microscopy, histology and immunostaining

Immunofluorescent analysis of BMDMs were performed seeding 1.5 × 10^4^ cells on sterile 12mm coverslips (Neuvitro Corporation, GG-12-1.5-Fibronectin). Cells were treated as indicated and then fixed with 4% paraformaldehyde. After permeabilization using 0.1% Triton X-100 0.2% BSA-PBS, cells were blocked in 2% BSA-PBS and incubated with NF-κB p65 (1:100, Cell Signaling Technology, Cat# 6956S) followed by Alexa Fluor 488 conjugated chicken anti-mouse IgG (1:300, Thermo Fisher Scientific, Cat# A-21200). Nuclei were stained with DAPI (1:10000, Thermo Fisher Scientific, Cat# D3571).

Histological analysis on collected spleens and lungs were performed at Histology core facility and Confocal imaging and Immunohistochemistry (IHC) core facility at Beth Israel Deaconess Medical Center. The IHC-DAB staining was performed on paraffin embedded mouse spleen sections. Slides were cut, deparaffinized and hydrated. Antigen retrieval was performed by boiling the slides in 10 mM sodium citrate (pH 6) in a pressure cooker. Endogenous peroxidase activity was blocked with 3% hydrogen peroxide and sections were blocked with 2% BSA (Jackson ImmunoResearch Lab, Cat# 001-000-162) for 1h at room temperature. After serum blocking, the slides were incubated with rabbit anti-oxidized phospholipid E06 IgG (1:100, Absolute Antibody, Cat# Ab02746-23.0) primary antibody overnight at 4°C. After wash, the slides were incubated with goat anti-rabbit IgG conjugated with HRP polymer (Abcam, Cat#: ab214880) for 90 minutes at room temperature. The slides were washed before developing in DAB (diaminobenzidine) metal enhanced kit (Vector lab, Cat# SK4105) and counter stained with hematoxylin (Thermo Fisher Scientific, Cat# 411160250). Lastly, the IHC slides were processed through dehydration steps before mounted in Permount (Fisher Scientific, Cat# SP15-100). The triple immunofluorescence antigen labeling was performed on paraffin-embedded mouse spleen sections. The paraffin sections were deparaffinized and rehydrated, followed by antigen retrieval using Tris-EDTA buffer (pH 9). After three washes, the sections were incubated with 2% BSA for 1 h at room temperature. Slides were then incubated with rabbit anti-oxidized phospholipid E06 IgG (1:100, Absolute antibody, Cat#: Ab02746-23.0), mouse anti-CD68 (1:100, Santa Cruz Biotechnology, Cat# sc-20060) and rat anti-Ly6G/Ly6C (1:50, Santa Cruz Biotechnology, Cat#: dc-59338) overnight at 4°C. The slides were washed three times and incubated with Alexa Fluor 488 conjugated donkey anti-rabbit (1:300, Jackson ImmunoResearch Lab, Cat# 711-545-152), Cy3 conjugated donkey anti-mouse secondary antibody (1:300, Jackson ImmunoResearch Lab, Cat# 715-165-151) and Alexa Fluor 647 conjugated donkey anti-rat secondary antibody (1:300, Jackson ImmunoResearch Lab, Cat# 712-605-153) for 90 minutes at room temperature. Samples were counterstained with Hoechst 33342 (1:10000, Life Technology, Cat#: H21492), washed and then mounted in Prolong Gold anti-fade mounting media (Thermo Fisher Scientific, Cat#: P36930). To analyze tissue inflammation and damage in lungs, the organs were collected, fixed in 10% formalin and processed for paraffin inclusion. 5 μm sections were stained with hematoxylin and eosin and perivascular and peribronchial inflammation were evaluated by severity-based ordinal scoring, based on cellular infiltration: 0 = none, 1 = solitary infiltration of cells, 2 = a ring of inflammatory cells 1 cell layer deep, 3 = a ring of inflammatory cells 2–4 cells deep and 4 = a ring of inflammatory cells >4 cells deep. Interstitial inflammation score was evaluated as follow: 0 = none, 1 = presence of few cells in septa, 2 = increased alveolar thickness and moderate cellular infiltration of septa, 3 = severe cellular infiltration in thickened septa and in air-space lumens. Each score was normalized for the surface of affected lung. The final inflammation score of each lung (0-11) was obtained by the sum of the normalized scores of perivascular, peribronchial and interstitial inflammation.

All the images were acquired using EVOS M7000 Imaging System (Thermo Fisher Scientific, Cat# AMF7000) and analyzed using ImageJ (NIH).

### In vivo models

The animals were housed in the same vivarium room at the Boston Children’s Hospital animal facility, up to five animals per cage, under controlled temperature, 12:12 light-dark cycle, and free access to normal food and water. All animal experiments were performed under the Guide of Care and Use of Laboratory Animals of the National Institute of Health. All animal experiments were approved by the Institutional Animal Care and Use Committee (IACUC) of Boston Children’s Hospital.

Indicated mice (females, 8-10 weeks old) were kept under specific-pathogen-free conditions and were co-housed for at least 4 weeks before any experiment.

For PRR challenge, PRR agonists (LPS, CpG, R848 or WPG) were i.p. injected, at the indicated concentrations, and blood and spleens were collected at 8h and 24h.

In CLP experiments, the day before the procedure, mice were implanted with temperature transponders (Bio Medic Data Systems, Cat# IPTT-300) and the abdomen was shaved. For acute IL-10 signaling inhibition, the following antibodies were i.p. injected at 0.5 mg per mouse: rat anti-mouse CD210 (IL-10R, clone 1B1.3a, BioLegend, Cat# 112712, Lot# B360179,) and rat IgG1, isotype control (BioLegend, Cat# 400458, Lot# B302875). For EZH2 inhibition, GSK343 (10 mg Kg^−1^), resuspended in 10% DMSO, 10% PEG-400, 10% Tween-80, 70% saline, was i.p. injected 48h and 12h before the experiment. For CLP procedure, mice were anesthetized, and cecum was exteriorized. The cecum was tightly suture ligated at about 1 cm to the distal end and was perforated by single through-and-through puncture midway between the ligation and the tip of the cecum in a mesenteric-to-antimesenteric direction with a 20G needle. After removing the needle, the cecum was then gently squeezed to extrude a small amount of feces. The cecum was relocated into the abdominal cavity, the peritoneum was closed by applying simple running sutures and skin was closed using metallic clips. In sham group, mice were operated following the same protocol without CLP procedure. After operation, mice were resuscitated by subcutaneously injecting pre-warmed saline and the analgesic buprenorphine extended-release (Fidelis Animal Health, Ethiqa XR) was administered subcutaneously at 3.25 mg Kg^−1^. Animals were monitored for 4 days for body temperature drop (Bio Medic Data Systems, Bluetooth Wireless Reader, Cat# DAS-8027-BLU System), weight loss and mortality comparison. Blood and spleens were collected at the indicated time points. Blood was left undisturbed at room temperature for 15 minutes and centrifugated at 4°C 2000 x g for 10 minutes to collect serum. Serum was immediately used for the measurement of cytokines and oxPLs levels. Alternatively, serum was immediately stored at −80°C.

The MRSA experiments were conducted at the University of California San Diego (UCSD) and protocols were approved by the Institutional Animal Care and Use Committee (IACUC) of UCSD. Male and female mice were injected with MRSA via tail vein with a lethal dose (1.8 x 108 CFU/mouse) and monitored for mortality over 120 hours. Blood was collected at 24 hours for measurement of bacterial load. In a separate experiment, male mice were injected with a sublethal dose (1.0 x 108 CFU/mouse) and monitored for 120 hours for mortality. Serum was collected for cytokine quantification at 24 hours.

To mimic persistent or post-acute viral infection, 2.5 mg Kg^−1^ of poly(I:C) or saline were daily administrated in mice via intratracheal instillation for the indicated time points. Body temperature, recorded using subcutaneous transponders (Bio Medic Data Systems, Cat# IPTT-300), and body weight were monitored every 24 hours. Survival was followed overtime, using humane endpoint at weight loss >20%. After 6 days, mice were euthanized by CO_2_ inhalation. Mice were transcardially perfused with 10 ml of PBS or until liver and lungs were clear. Lungs were harvested and processed for histology inspection or lysed in RIPA buffer, added with protease and phosphatase inhibitors, using a Bead Mill 24 Homogenizer (Fisher, Cat# 15-340-163). After centrifugation at 17000 x g for 10 minutes at 4°C, the resulted clear tissue homogenates were immediately used for the quantification of cytokines and oxPLs.

### Bacterial Counts

Collected whole blood was serially diluted in sterile PBS and cultured overnight at 37°C on 5% blood agar plates (Hardy Diagnostics, Cat# A10BX). Bacteria colonies were counted, and the results were expressed as log of colony-forming units (CFU) per ml.

### Metabolomics analysis

Metabolomics profiling was performed at the Beth Israel Deaconess Medical Center Mass Spectrometry Facility. Briefly, polar metabolites were collected by methanol-based extraction, as previously described [58]. Samples were resuspended using HPLC-grade water for mass spectrometry and analyzed using a 5500 QTRAP hybrid triple quadrupole mass spectrometer (AB Sciex) coupled to a Prominence UFLC HPLC system. A total of 284 endogenous water-soluble metabolites were analyzed). Statistical analysis was performed using MetaboAnalyst (http://www.metaboanalyst.ca, free online software).

### Microscale thermophoresis (MST)

Microscale thermophoresis experiments were performed at the Harvard Center for Macromolecular Interactions (CMI). His-tagged recombinant human AKT1 (BPS Bioscience, Cat# 40003), AKT2 (BPS Bioscience, Cat# 40011) and AKT3 (BPS Bioscience, Cat# 40012) were fluorescently labeled using His-Tag Labeling Kit RED-Tris-NTA 2^nd^ Generation (NanoTemper, Cat# MO-L018). The proteins (at fixed concentration) were incubated with oxPAPC or DPPC, at the indicated concentrations, and samples were loaded in Monolith capillaries (NanoTemper, Cat# MO-K022). MST was run in Monolith NT.115 Instrument (NanoTemper) and analyzed with MO Affinity Analysis software, Version 2.3 (NanoTemper).

### Kinase activity

The inhibition of kinase activity mediated by phospholipids was determined using the following Z’-LYTE Kinase Assay Kits: Z’-LYTE Kinase Assay Kit - Ser/Thr 6 Peptide (Thermo Fisher Scientific, Cat# PV3179) used to measure AKT activity and the Z’-LYTE Kinase Assay Kit - Ser/Thr 7 Peptide (Thermo Fisher Scientific, Cat# PV3180) used to measure PDK1 activity. All assays were run in accordance with the manufacturer’s instructions. Briefly, recombinant human AKT1 (BPS Bioscience, Cat# 40003), AKT2 (BPS Bioscience, Cat# 40011) and AKT3 (BPS Bioscience, Cat# 40012) were used at 7.5 ng per reaction and recombinant human PDK1 (BPS Bioscience, Cat# 40080) was used at 50 ng per reaction. Phospholipids were prepared by 2-fold serial dilution in kinase buffer and incubated with the target protein for at least 10 minutes at room temperature. Specific Fluorescence Resonance Energy Transfer (FRET)-peptides were added at 2 μM and the solution was supplemented with ATP (100 μM for AKT or 50 μM for PDK1) to initiate the reaction.

ATP was omitted in control reactions to obtain a complete kinase activity inhibition (100% inhibition). After 1h, development reagent, that selectively cleaves non-phosphorylated peptides disrupting FRET between the donor (coumarin) and acceptor (fluorescein) fluorophores on the FRET-peptides, was added and incubated for 1 hour at room temperature. After stopping the reaction with stop reagent, coumarin and FRET based fluorescein emission was measured on a SpectraMax i3x plate reader (Molecular Devices).

### Immunoprecipitation

293T cells were transiently transfected using Lipofectamine 3000 (Thermo Fisher Scientific, Cat# L3000001) with the following plasmids: pCDNA3-HA-Akt1 (Addgene plasmid # 73408, RRID:Addgene_73408), pCDNA3-HA-Akt1-aa1-149 (Addgene plasmid # 73410, RRID:Addgene_73410), pCDNA3-HA-Akt1-aa1-408 (Addgene plasmid # 73412, RRID:Addgene_73412) and pCDNA3-HA-Akt1-aa120-433 (Addgene plasmid # 73411, RRID:Addgene_73411). These plasmids were a gift from Jie Chen (University of Illinois at Urbana-Champaign) [59]. Cell lysates were incubated with oxPAPE-N-Biotin (10 μg) with or without competing oxPAPC doses at 4°C on a nutator for either 6h or overnight. Lipid-protein complexes were captured using High Capacity Streptavidin Agarose beads (Thermo Fisher Scientific, Cat# 20357) and then heated at 98°C in Laemmli Sample Buffer (Bio-rad, Cat# 1610747). Samples were run in a SDS-PAGE gel and lipid-interacting proteins were detected by immunoblotting using HA-tag antibody (Proteintech Group, Cat# 51064-2-AP).

### RNA-Seq

BMDMs (1 × 10^6^) were treated as indicated and RNA was purified using Direct-zol RNA Miniprep (Zymo Research Corporation, Cat# R2053). Samples were submitted to Azenta Life Sciences for standard RNA-seq processing. RNA was quantified using Qubit 2.0 Fluorometer (Thermo Fisher Scientific) and RNA integrity was checked with RNA Screen Tape on Agilent 2200 TapeStation (Agilent Technologies). After library preparation (Illumina, PolyA selection), samples were sequenced using Illumina, 2×150 bp, ∼350M PE reads (∼105GB), single index.

### ChIP-qPCR

BMDMs (5 × 10^6^) were treated as indicated and fixed with 1% formaldehyde for 10 minutes at room temperature. After quenching with 650 mM glycine for 5 minutes, cells were washed in chilled PBS and collected by scraping. Subsequent sample processing was performed using Chromatrap Enzymatic kit (Chromatrap Porvair Sciences, Cat# 500191). Briefly, cells were lysed, and chromatin was sheared by enzymatic digestion to obtain 150-500 bp fragments. Next, chromatin was incubated with H3K27me2me3 antibody (2μg, Active Motif, Cat# 39435) or mouse IgG isotype antibody (2μg, Santa Cruz Biotechnology, Cat# sc-2025) for 1h at 4°C and antibody-chromatin complexes were selectively collected by protein A spin columns. Samples were reverse cross-linked, and proteins digested with proteinase K. DNA fragments were purified using ChIP DNA Clean & Concentrator columns (Zymo Research Corporation, Cat# D5205).

Specific primer pairs were used to amplified conserved noncoding sequences (CNS) related to murine *Il10* locus. The following primers were obtained from Neumann et al. 2014 [60]: CNS −4.5 (Fwd: GCCACGATTCTCAGGACATT, Rev: GTATCCAACCCCACTTGCAC) and CNS −0.5 (Fwd: CTCTCCTCTGACCAACTGCC, Rev: TGGGTTGAACGTCCGATATT). The following primers were obtained from Lee et al. 2012 [61]: CNS −0.12 (Fwd: TCTGTACATAGAACAGCTGTC, Rev: CTGGTCGGAATGAACTTCTG), CNS +1.65 (Fwd: GTCTCTTGCTCATCTGTCTC, Rev: GCTAATAACCCACAATGACTC), CNS +2.98 (Fwd: ACTAGGTGTTGAGGAGAGTG, Rev: GAATTCTGCTTTCTGCTCGT), CNS +6.45 (Fwd: GTGGTCATTTTTTCAGTAAGACC, Rev: CCTAACCTTTCATCTCACAG). Control primers were purchased from Active Motif: *Hoxc10* (Cat# 71019) and *Gapdh* (Cat# 71014).

### Statistical analysis

The statistical tests used and *P* values for all the experiments are specified in figure captions. Statistical significance for the experiments with more than two groups and two factors was tested with two-way analysis of variance (ANOVA), and Sidak’s, Dunnet’s or Tukey’s multiple-comparisons tests were performed according to the nature of the comparison tested. When comparisons between only two groups were made, an unpaired two-tailed *t*-test was used to assess statistical significance. For survival experiments, *P* values were determined by long-rank test. To establish the appropriate test, normal distribution and variance similarity were assessed with the D’Agostino–Pearson omnibus normality test. All statistical analyses were performed using Prism 9 (GraphPad Software, version 9.4.0).

